# IRF5 promotes intestinal inflammation by guiding monocyte differentiation towards a pathogenic CD11c^+^ macrophage phenotype

**DOI:** 10.1101/601963

**Authors:** Alastair L. Corbin, Maria Gomez-Vazquez, Tariq E. Khoyratty, Dorothée L. Berthold, Hannah Almuttaqi, Moustafa Attar, Isabelle C. Arnold, Fiona M. Powrie, Stephen N. Sansom, Irina A. Udalova

**Affiliations:** The Kennedy Institute of Rheumatology, University of Oxford, Oxford. UK; Institut für Molekulare Krebsforschung, University of Zurich, Zurich, Switzerland

## Abstract

Mononuclear phagocytes (MNPs) play a key role in maintaining intestinal homeostasis but also in triggering immunopathology in response to acute microbial stimulation, which induces the recruitment of masses of Ly6C^hi^ monocytes to the gut. The regulators that control monocyte tissue adaptation in the gut remain poorly understood. Interferon Regulatory Factor 5 (IRF5) is a transcription factor previously shown to play a key role in maintaining the inflammatory phenotype of macrophages. Here we investigate the impact of IRF5 on the MNP system and physiology of the gut at homeostasis and during inflammation. We demonstrate that IRF5 deficiency has a limited impact on colon physiology at steady state, but ameliorates immunopathology during *Helicobacter hepaticus* induced colitis. Inhibition of IRF5 activity in MNPs phenocopies global IRF5 deficiency. Using a combination of bone marrow chimera and single cell RNA-sequencing approaches we compare the differentiation trajectories of wild type and IRF5 deficient monocytes in a shared inflammatory environment and demonstrate that IRF5 stipulates a choice in monocyte differentiation towards macrophages. Specifically, IRF5 promotes the generation of pathogenic CD11c^+^ macrophages and controls the production of inflammatory mediators by these cells. Thus, we identify IRF5 as a key transcriptional controller of pathogenic monocyte differentiation in the gut.

## Introduction

The term Inflammatory Bowel Disease (IBD) encompasses a group of debilitating inflammatory conditions of the gastrointestinal tract that affects ∼0.5-1% of westernised populations^1^. The IBDs are associated with high morbidity and burden healthcare systems^2, 3^. Conventional IBD therapies are limited by moderate-high rates of adverse events, or patient unresponsiveness, whilst approximately 40% of patients successfully treated with anti-TNFα become refractory to therapy^2^. Therefore, there is unmet clinical need for IBD therapies. The aetiology of IBD is unknown, but interplay between host genetics, and environmental factors, and the microbiota contribute to disease pathogenesis^1^.

Mononuclear Phagocytes (MNPs), including monocytes, macrophages, and Dendritic Cells (DCs), are present in large numbers in the colonic Lamina Propria (cLP), and carry out diverse, overlapping functions critical to the maintenance of intestinal homeostasis. The dysregulation of the intestinal MNP system leads to infection and inflammation^4 5, 6, 7, 8, 9, 10, 11^.

The origins of the intestinal MNP systems has been the topic of considerable debate in recent years, clouded by inconsistent nomenclature and shared surface markers between macrophages and DCs. Intestinal Lamina Propria DCs at the steady state are largely derived from pre-DC precursors, generated in the bone marrow, which are understood differentiate into three major intestinal DC subsets. These subsets comprise of an XCR1 positive (Xcr1^+^SIRPα^−^CD103^+^Cd11b^−^CX3CR1^−^) population that is analogous to classical dendritic cells (cDC) 1 DCs, and two cDC2-like SIRPα positive (SIRPα^+^Xcr1^−^Cd11b^+^CX3XR1^+^) subsets which can be further discriminated by CD103 expression^12, 13^. In addition, the existence of a discrete population of hybrid macrophage/DC cells within the cDC2 intestinal compartment has been described^14^. The ontogeny of intestinal DCs during inflammation is more complicated since some monocyte-derived cells may acquire phenotypic and functional DC hallmarks^15, 16, 17^.

Intestinal Lamina Propria macrophages have dual origins: from embryonically derived macrophages (CD4^+^ Tim4^+^) that self-renew, and monocytes, but in the adult mouse, most of the macrophage turnover is of monocytic origin^18, 19^. In mice, the differentiation of monocytes to macrophages in the cLP has been termed the “monocyte waterfall” ^19^. After entering the cLP, naïve Ly6C^hi^, MCHII^−^ (P1) monocytes begin maturing by acquiring expression of MHCII (P2) before downregulating Ly6C expression. The pool of MHCII^+^ cells comprises of Ly6C^+/−^CX3CR1^Int^ monocyte/macrophage intermediates (P3) and fully mature Ly6C^−^CX3CR1^hi^F4/80^hi^CD64^hi^MHCII^hi^ macrophages (P4) ^19, 20^. The blood origin of intestinal macrophage subsets was also confirmed in human studies where two monocyte-derived macrophage populations: CD11c^+^ with high turnover and CD11c^−^ with slow turnover, were identified at steady state^21^. It was suggested that CD11c^+^ macrophages might be an intermediate between blood monocytes and tissue resident CD11c^−^ macrophages^21^.

The regulators that control the transition of monocytes through a number of intermediate differentiation states are largely unknown, but the cytokines, TGFβ and IL10, have been linked to the development of cLP tissue-resident macrophages^22, 23^. CX_3_CR1^IL10R−^ mice exhibited heightened inflammation, which maintained a pro-inflammatory mono-macrophage state, preventing their full differentiation and initiating spontaneous colitis ^23^. Loss of TGFβ-Receptor on macrophages resulted in a minor impairment of macrophage differentiation, defined by transcriptional profiling of monocyte to macrophage transition in the cLP ^22^.

One candidate intrinsic regulator of the intestinal macrophage signature is Interferon Regulatory Factor 5 (IRF5), which was described to promote an inflammatory macrophage phenotype^24^ and has variants that are genetic risk factors for Ulcerative Colitis and Crohn’s Disease^25 26 27^. IRF5 is activated by phosphorylation and ubiquitination events downstream of Pattern Recognition Receptors (PRRs), e.g. NOD2, TLR2, and TLR4, and directly regulates many cytokines associated with IBD (IL-1β, IL-6, IL-10, IL-12, IL-23, TNF), placing IRF5 as a nexus for the regulation of inflammatory responses^1, 24, 28^. To formally examine the role of IRF5 in the establishment of intestinal MNP system, we compared the continuum of cell states of wild type (WT) and IRF5 deficient (*Irf5*^−/−^) MNPs at homeostasis and during *Helicobacter hepaticus* (*Hh*) induced intestinal inflammation using a combination of competitive Mixed Bone Marrow Chimaera (MBMC), single cell gene expression analysis (scRNA-seq) and functional validation approaches. Hh infection concomitant with the administration of anti-Interleukin 10 Receptor (αIL10R) antibodies triggers IL-23 dependent intestinal inflammation with robust T_H_1/T_H_17 T cell response, which carries many features of human IBD^9, 29, 30, 31^. In this model, CX_3_CR^int^ and CD11c^+^ monocyte/macrophages intermediates drive immunopathology by producing pro-inflammatory cytokines such as IL-23, IL-1β and TNFα^9 32^. We report a novel function for IRF5 in controlling the cell fate choice of Ly6C^hi^ monocytes in the tissue, delineating the pathogenic CD11c^+^ macrophages in inflammation and controlling the immunopathology of Hh + aIL10R-induced colitis.

## Results

### IRF5 deficiency has limited impact on colon physiology at steady-state

IRF5 expression in the cLP has not been described, nor the effects of IRF5 deficiency. Colons (Fig.1A), and caeca (Supplementary Fig.S1A) of WT and *Irf5*^−/−^ were comparable in morphology. Sections were scored for epithelial hyperplasia, nucleated cell infiltrate, area affected, and submucosal oedema and displayed no obvious signs of inflammation (score < 3) and no morphological differences between WT and *Irf5*^−/−^ (Fig. 1B **and** Supplementary Fig S1B). The immune compartment of the cLP was evaluated by flow cytometry and revealed that the number of leukocytes in the colon (live CD45^+^) were comparable between WT and *Irf5*^−/−^ (Fig. 1C).

**Figure 1:**
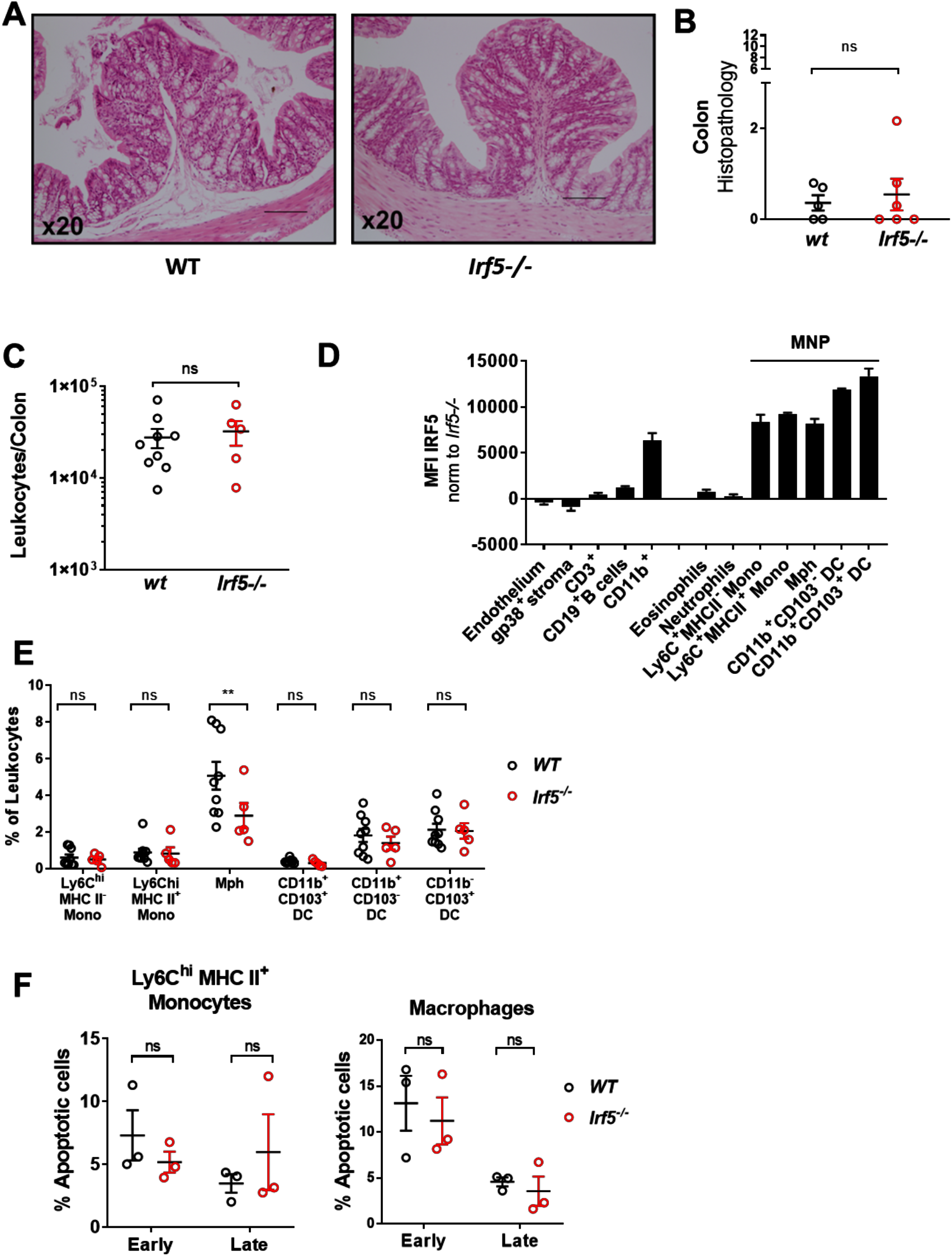
*Irf5^−/−^ mice* display reduced frequencies of macrophages in the colonic lamina propria. **A)** Representative H&E sections of colons from WT (left) and *Irf5*^−/−^ (right) mice at steady state. **B)** Histopathology scoring of WT (n=6) and *Irf5*^−/−^ (n=5) colons. **C)** Number of cLPLs retrieved from steady state WT (n=9) and *Irf5*^−/−^ (n=5) mice. **D)** IRF5 expression in the steady state cLP of WT mice (n=3). **E)** The frequency of intestinal MNPs in the cLP of steady state WT (n=9) and *Irf5*^−/−^ (n=5) mice. **F)** Quantification of early phase (Annexin V^+^ Live/Dead^−^), and late phase (Annexin V^+^ Live/Dead^+^) cell death quantified by Annexin V staining in WT (n=3) and *Irf5*^−/−^ (n=3) cLP Ly6C^hi^ MHC II^+^ (P2) monocytes and macrophages. **B, C, E:** Data are pooled from two independent experiments. **D, F:** data are representative of two independent experiments. **B, C)** Mann-Whitney U test. **E, F)** Two-Way ANOVA with Sidak correction. Data presented are mean ± SEM, ns = not significant, ** p ≤ 0.01

Next, we assessed the levels of IRF5 expression in the cells of the colon and demonstrated that non-myeloid, and non-leukocyte populations expressed low levels of IRF5 compared to CD11b^+^ myeloid cells (Fig.1D). Among myeloid cells, MNPs, i.e. monocytes, macrophages and DCs, expressed the highest levels of IRF5 (Fig.1D). The composition of the cLP myeloid compartment in WT and *Irf5*^−/−^ was profiled using the gating strategy that included definition of the stages of monocyte differentiation^9, 20^ (Supplementary Fig.S1D). Frequencies of Ly6C^hi^MHCII^−^ (P1) and Ly6C^hi^MHCII^+^ (P2) monocytes and CD11b^+^ DCs among the infiltrated leukocytes were similar in *Irf5*^−/−^ animals but a higher frequency of fully differentiated F4/80^+^ macrophages was observed in WT mice (5.1%) than in *Irf5*^−/−^ (2.9%) (Fig.1E). IRF5-deficient Ly6C^hi^ MHC II^+^ monocytes and macrophages were no more sensitive to apoptosis than their WT counterparts (Fig. 1F). Thus, we hypothesised that IRF5 may promote differentiation of monocytes to macrophages.

### IRF5 deficiency protects against intestinal inflammation

Next, we evaluated the effect of IRF5 deficiency on the pathogenesis of intestinal inflammation. The commensal bacterium, Hh, is commonly found in the mucosal layer of the murine lower intestinal track, including colon and caecum, and does not lead to inflammation in WT animals. However, the concomitant administration of anti-IL10receptor (αIL10R) monoclonal antibodies in Hh-naïve mice triggers inflammation, with animals developing histological and molecular signatures of colitis within 2-3 weeks post infection^9^. WT and *Irf5*^−/−^ mice were subjected to Hh + αIL10R colitis for 21 days and inflammatory indices were analysed upon sacrifice. Morphological analysis (Fig 2A, Supplementary Fig. S2A) and histological scoring indicated that both colons (Fig. 2B) and caeca (Supplementary Fig. S2B) were protected from colitis by IRF5-deficiency. The leukocyte infiltrate to the cLP was significantly reduced in *Irf5*^−/−^ mice (Fig. 2C), consistent with the reduced levels of inflammation in the colon and caecum of *Irf5*^−/−^ animals. Next, we profiled T_H_1 and T_H_17 lymphocyte responses that are involved in the pathogenesis of colitis^33^. *Irf5*^−/−^ mice displayed a significantly reduced T_H_1 effector response as quantified by emergence of IFNγ^+^ CD4^+^ T cells, and a non-significant reduction in the number of T_H_17 cells (Fig. 2D).

**Figure 2:**
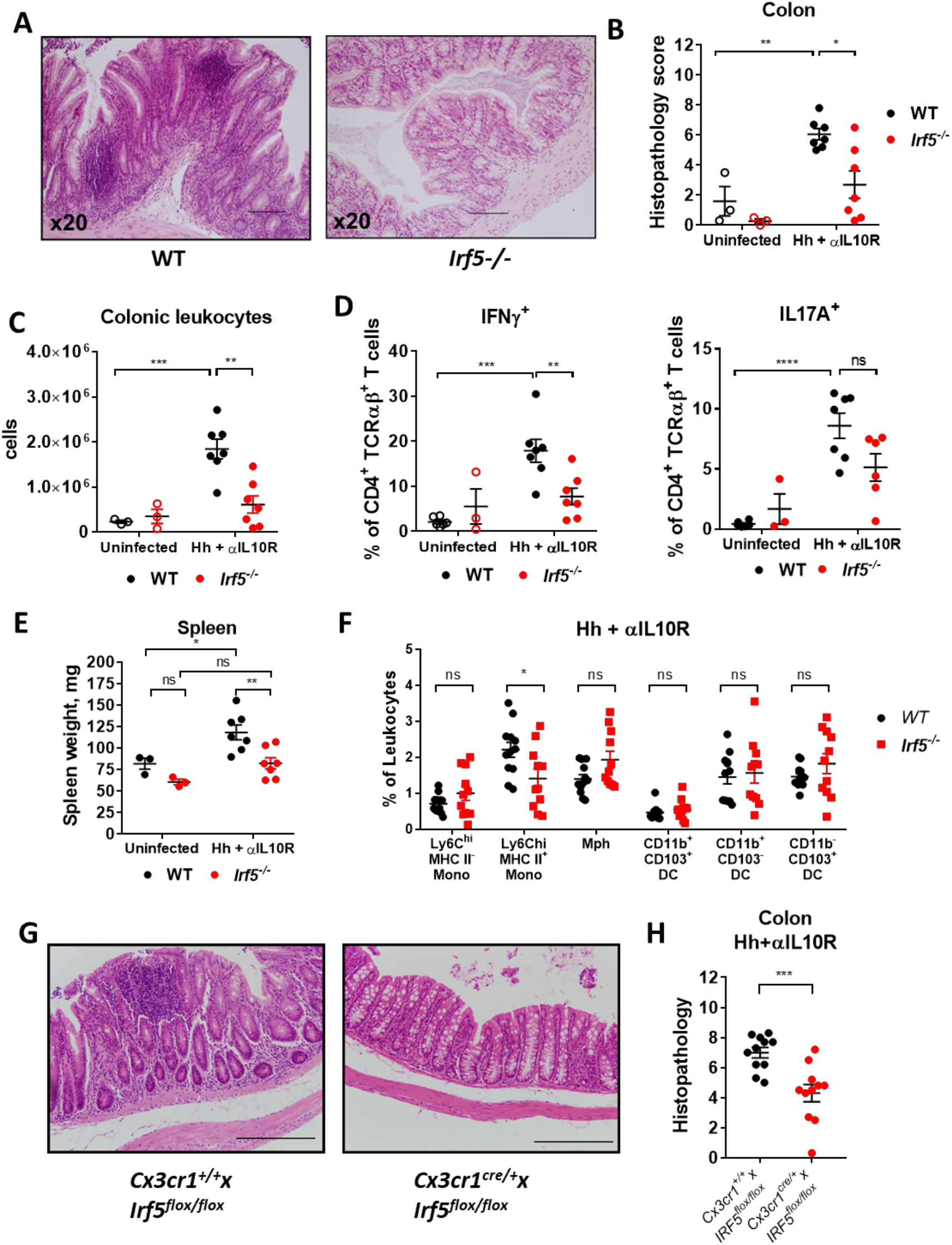
IRF5 promotes intestinal inflammation in response to *Helicobacter hepaticus* + αIL10R. **A)** Representative H&E sections of colons from WT (left) and *Irf5*^−/−^ (right) mice at d21 Hh + αIL10R. **B)** Histopathology scoring of WT and *Irf5*^−/−^ colons. **C)** Number of cLPLs retrieved from steady state and d21 Hh + αIL10R WT and *Irf5*^−/−^ mice. **D)** frequencies of IFNγ-producing and IL17A-producing CD4^+^ T cells in WT and *Irf5*^−/−^ mice. **E)** Spleen weights of WT and *Irf5*^−/−^ mice at steady state and d21 Hh + αIL10R. **F)** The frequency of intestinal MNPs in the cLP of steady state WT (n=12) and *Irf5*^−/−^ (n=11) mice. **G)** Representative H&E sections of colons from CX_3_CR1^IRF5+^ (left) and CX_3_CR1^IRF^^5−^ (right) mice at d21 Hh + αIL10R. **H)** Histopathology scoring of colons from CX_3_CR1^IRF5+^ (n=11) and CX_3_CR1^IRF^^5−^ (n=11) mice at d21 Hh + αIL10R. **B, C, D, E:** Data are representative of two independent experiments, (ss n=3, Hh + αIL10R n=7). **F,H:** Data are pooled from two independent experiments. **B, C, D, E)** Two-Way ANOVA with Tukey correction, **F)** Two-Way ANOVA with Sidak correction, **H)** Unpaired t-test. Data presented are mean ± SEM, ns = not significant, * p ≤ 0.05, ** p ≤ 0.01, *** p ≤ 0.001, **** p < 0.0001

Hh + αIL10R colitis in WT mice led to splenomegaly, but not in *Irf5*^−/−^ mice (Fig. 2E), indicating that they were also protected from systemic aspects of disease. Despite the altered immune response, Hh presence in the caecal faeces were unaffected in *Irf5*^−/−^ compared to WT, quantified by detection of the Hh Cytolethal Distending Toxin B (cdtB) gene (Supplementary Fig. S2C), ruling out differential bacterial colonisation in WT and IRF5-deficient animals.

Myeloid cells make up a significant part of the leukocyte pool at the peak of inflammation in the colon^32^. Masses of Ly6C^hi^ monocytes are rapidly recruited to the gut in response to inflammatory signals, with Ly6C^hi^MHCII^+^ inflammatory monocytes becoming the predominant cells that carry out inflammatory effector functions^9, 15, 16, 20, 23, 34, 35^. Indeed, we observed a significant increase in Ly6C^hi^MHCII^+^ inflammatory monocytes at the peak of Hh + αIL10R induced inflammation, while the frequency of F4/80^+^ macrophages diminished (Supplementary Fig. S2D). The frequencies of the DC populations remained unaffected by ongoing inflammation (Supplementary Fig. S2D). IRF5 deficiency attenuated the predominance of Ly6C^hi^MHCII^+^ inflammatory monocytes (Fig. 2F), approximating the monocyte-macrophage waterfall observed at the steady state (Fig. 1C). Thus, we concluded that the curtailed inflammatory environment of the IRF5-deficient animals imprints on the waterfall of monocyte differentiation re-directing it towards the steady state.

To confirm that IRF5 activity in MNPs is a major contributor into the immunopathology of intestinal inflammation, we subjected Cx3cr1^cre^xIRF5^flox/flox^ animals, which are deficient in IRF5 specifically in their MNP compartment (Supplementary Fig S2E), to the Hh + αIL10R colitis model. The inflammatory indices were analysed upon sacrifice of mice. Histological scoring indicated that both colons (Fig. 2G, H) and caeca (Supplementary Fig S2F, E) were protected from colitis by IRF5-deficiency in MNPs.

These data demonstrate that IRF5 plays a critical role in the pathogenesis of intestinal inflammation largely acting via the MNP system.

### IRF5 promotes generation of macrophages in cell-intrinsic manner

At homeostasis, a higher frequency of fully differentiated F4/80^+^ macrophages was observed in WT compared to *Irf5*^−/−^ mice (Fig 1), suggesting that IRF5 may play role in monocyte differentiation. However, in Hh + aIL10R induced colitis, the different inflammatory environments between WT and *Irf5*^−/−^ animals obscured this effect (Fig 2). To compare the differentiation competence of WT and *Irf5*^−/−^ monocytes in a shared environment, we performed mixed bone marrow chimera (MBMC) experiments. The lethally irradiated mice were reconstituted with 50:50 WT*:Irf5*^−/−^ bone marrow mix and the efficiency of reconstitution in the bone marrow, of blood monocytes, and of the cLP MNP compartment were investigated. We observed no difference in reconstitution of long term (LT)- or short term (ST)-haematopoietic stem cells (HSCs), myeloid progenitors (common myeloid progenitors (CMPs), granulocyte-monocyte progenitors (GMP) and megakaryocyte-erythrocyte progenitors (MEP) or Ly6C^hi^ mature monocyte population in the bone marrow (Supplementary Fig S3A, B). IRF5 expression assessed by intracellular staining using flow cytometry was negligible in LT-HSCs, ST-HSCs, and CMPs but detectable in GMPs and MEPs. Ly6C^hi^ monocytes express the highest levels of IRF5 among the tested progenitor and mature cell populations (Supplementary Fig. S3C). The reconstitution of Ly6C^hi^ monocytes in the blood was not affected by IRF5 deficiency, but Ly6C^hi^ monocytes in the cLP were preferentially derived from WT progenitors (Supplementary Fig. S3D, p < 0.0001). As progenitor cells were depleted by lethal irradiation of mice, and IRF5-deficient monocytes were not more sensitive to apoptosis (Supplementary Fig. S3E), we concluded that the recruitment of donor WT monocytes was more efficient than the recruitment of donor *Irf5*^−/−^ monocytes, consistent with previously published analysis^36^. WT monocytes also more readily migrated towards CCL2, a key monocyte chemoattractant in the cLP^20^ in an *in vitro* Boyden chamber analysis of the monocyte chemotaxis (Supplementary Fig. S3F).

When the composition of MNP pools was analysed, no difference in the frequencies of WT and *Irf5*^−/−^ Ly6C^hi^MHCII^−^ (P1) and Ly6C^hi^MHCII^+^ (P2) monocytes was observed in the MBMCs (Fig. 3A). However, consistent with the global *Irf5*^−/−^ phenotype at steady state (Fig. 1A), the frequency of WT macrophages within the MNP pool was significantly higher than *Irf5*^−/−^ macrophages (30.4% vs 23.8%, p = 0.0017) (Fig. 3A). Moreover, when the influence of extrinsic factors was minimised by the shared environment of MBMCs, we observed a greater frequency of WT than *Irf5*^−/−^ macrophages (18.3% vs 14.5%, p = 0.0001) under the Hh + aIL10R induced inflammation as well (Fig. 3B), suggesting that IRF5 acts cell-intrinsically to support the generation of intestinal macrophages. Of interest, we also observed that the CD11b^+^ DCs were overrepresented in uninfected *Irf5^−/−^* MBMCs and in inflammation (Fig. 3A, B).

**Figure 3:**
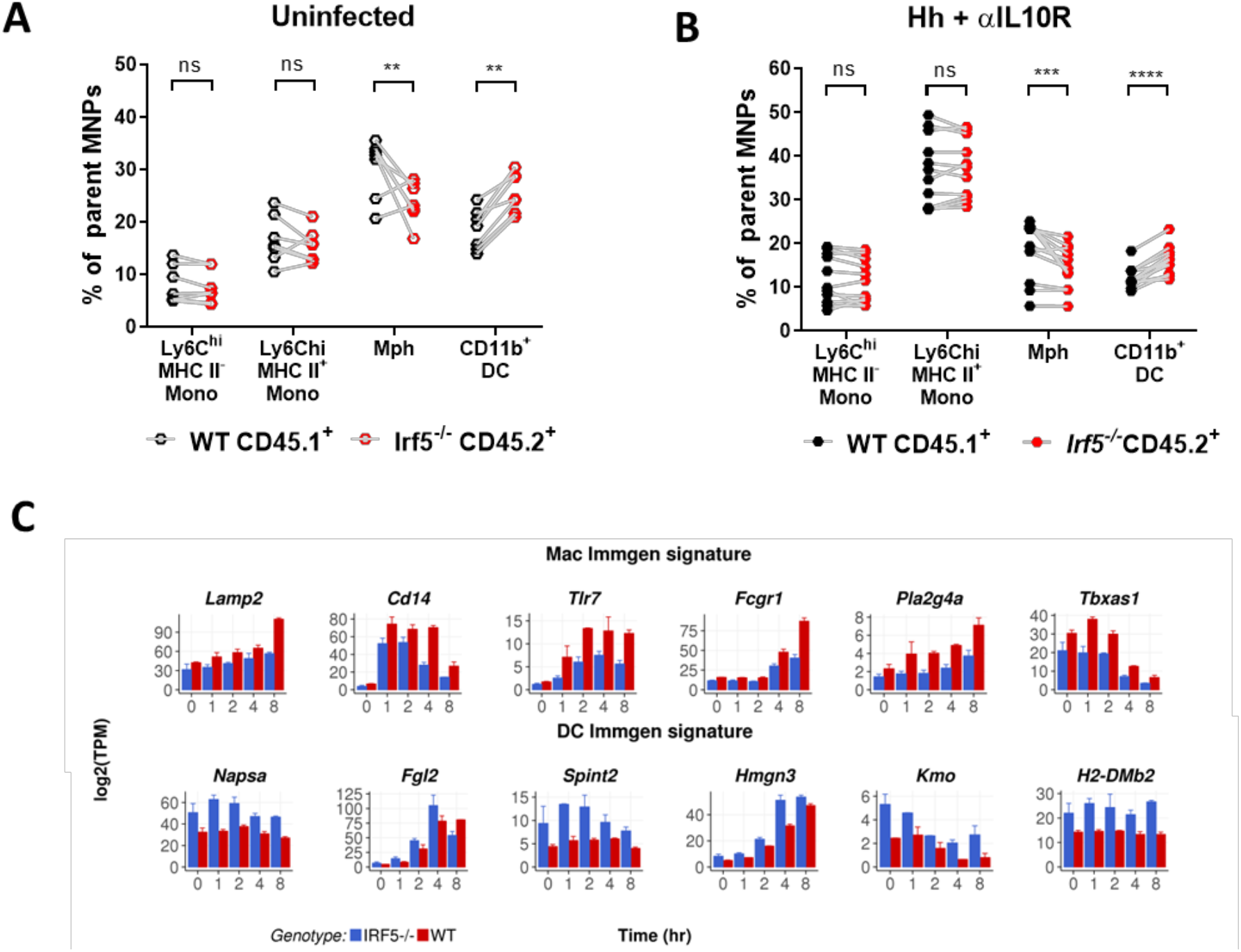
IRF5 deficiency precludes cLP monocytes from macrophage fate. The relative compositions of the MNP compartment of WT- and *Irf5*^−/−^-donor derived cLPLs were analysed by flow cytometry in **A)** Uninfected (n=7) and **B)** d21 Hh + αIL10R (n=12) MBMCs. **C)** Significantly differentially expressed Immgen signature genes between WT and *Irf5*^−/−^ GM-BMDMs stimulated with LPS. **A, B**: Data are pooled from two independent experiments. **C:** one experiment, n=2. **A, B)** Two-Way ANOVA with Sidak correction, **C)** RNAseq presented as log_2_TPMs (see materials and methods for statistics). Data presented are mean ± SEM, ns = not significant, ** p ≤ 0.01, *** p ≤ 0.001, **** p < 0.0001

To assess whether IRF5 might have a generic effect on mononuclear cell specification, we generated macrophages and DCs *in vitro* using bone marrow cultures supplemented with GM-CSF^37^. No significant effect of IRF5 deficiency on the generation of either cell type was detected in naive bone marrow cultures (Supplementary Fig. S3G). However, the analysis of known signatures of macrophage and dendritic cell gene expression^38, 39^ demonstrated that, following activation with LPS, *Irf5*^−/−^ cells upregulated DC specific-genes including *Zbtb46*, *Ciita*, *H2-DMb2* and *Kmo* whilst down-regulating genes associated with a macrophage identity including *Cd14*, *Fgcr3* (*Cd16*), *Fcgr4 (Cd64) Tlr7* and *Tlr4* (Fig 3c, Supplementary Fig. S3H). This suggests a role for IRF5 in controlling the expression of genes associated with cellular identity: macrophage associated genes up-regulated by LPS are more strongly expressed in WT cells than *Irf5*^−/−^, while a subset of LPS up-regulated DC genes are more highly expressed in the absence of IRF5. As the TLR4/Myd88 signalling cascade is important for the establishment of intestinal homeostasis^40^, we examined whether it might be modulated by IRF5, as previously shown for MAPK kinases in human myeloid cells^41^. No difference in phosphorylation of the MAP kinase, ERK, was detected in WT and *Irf5*^−/−^ GM-CSF derived cells in response to LPS (Supplementary Fig. S3H). Together, these *in vivo* and *in vitro* findings argue that, downstream of active PRR signalling, IRF5 biases monocytes toward a macrophage, rather than a dendritic cell-like fate.

### IRF5 defines an inflammatory MNP signature during colitis

To examine if IRF5 controls the expression of macrophage defining genes in intestinal macrophages, we conducted small bulk RNA-seq analysis of WT and *Irf5*^−/−^ Ly6C^hi^MHCII^−^ (P1) monocytes, Ly6C^hi^MHCII^+^ (P2) monocytes and Ly6C^−^MHCII^+^F4/80^+^ macrophages (P4) (n=100 cells/sample) from each of the inflamed colons of three Hh + αIL10R MBMC animals. First, we identified the genes that showed significant variation between the WT monocyte and macrophage samples. Hierarchical clustering of these genes revealed that the transcriptome of P2 monocytes overlaps with the profiles of both P1 monocytes and macrophages in line with the concept that they represent a transitional state of monocyte to macrophage differentiation (Supplementary Fig. S4A). We also detected high expression levels for genes previously shown to be associated with mature intestinal macrophages, such as MHC molecules (*H2-M2*), tetraspanins (*Cd72, Cd81*), complement molecules (*C1qa, C1qb, C1qc*), chemokines (*Ccl5, Ccl8*) and phagocytic and immunoactivating receptors (*Fcgr4, Fcer1g, Cd300e*) in the macrophage populations^22^. Next, we identified genes that were significantly (BH adjusted p < 0.05, |fold change| > 2) regulated by IRF5 in each of the P1 (n=607 genes), P2 (n=761 genes) and macrophage (n=977 genes) compartments (Supplementary Figure S4B). Amongst the differentially expressed genes, immature *Irf5*^−/−^ P1 monocytes showed significantly lower levels of *Smad2* and *Kdm3a* which respectively transduce and positively regulate TGF-B and Jak2/Stat3 signalling, pathways of known importance for monocyte maturation (Supplementary Fig. S4C). In line with this observation IRF5 deficient macrophages failed to down-regulate genes highly expressed in P1 and P2 monocytes including *Plac8*, *Cdkn2d* (*P19ink4d*) and *Irf1* and also showed significantly lower expression of the MHC class II molecule *H2-M2* (Supplementary Fig. S4C). At the same time, in macrophages, loss of IRF5 reduced expression of the key pro-inflammatory cytokines (*Il-12b*, *Ccl11*, *Tnfsf13b/BAFF*), expression of the immunoactivating receptor *Cd300e*, tetraspanins (*Cd81* and *Cd72*) as well as *IL-10*. A significant reduction in the expression of the key pro-inflammatory cytokine *Il-12b* was also observed in the P2 monocytes. These changes were accompanied by up and down-regulation of the epigenetic regulators *Hdac2* and *Hdac9* in IRF5 deficient macrophages. At the pathway level, geneset enrichment analysis of Gene Ontology (GO) Biological Process categories revealed that IRF5 broadly modulated inflammatory pathways including “leukocyte activation”, “response to interferon-gamma”, “response to bacterium” and “regulation of T-cell activation” in both monocytes (P1 & P2), and macrophages (BH adjusted p-value < 0.1, Fig. 4A). Genes regulated by IRF5 in the P2 monocyte compartment displayed a significant enrichment of genes involved in “regulation of interleukin-12 production”, “interleukin 6-secretion”, “interleukin-1 production”, while “response to interleukin-1”, “TNF superfamily cytokine production” was characteristic of macrophages (Fig. 4A). These pathways, and specifically production of IL-23, IL-1, and TNF, have been previously associated with colitis development and/or IBD^9, 28, 32^. In independent experiments, using flow cytometry, we confirmed that colonic WT macrophages in the MBMCs produced higher levels of cytokines TNFα and IL-1β cytokines than *Irf5*^−/−^ cells (Fig. 4B, C). The surface expression of MHCII was higher on WT macrophages relative to *Irf5*^−/−^ (Fig. 4D).

**Figure 4:**
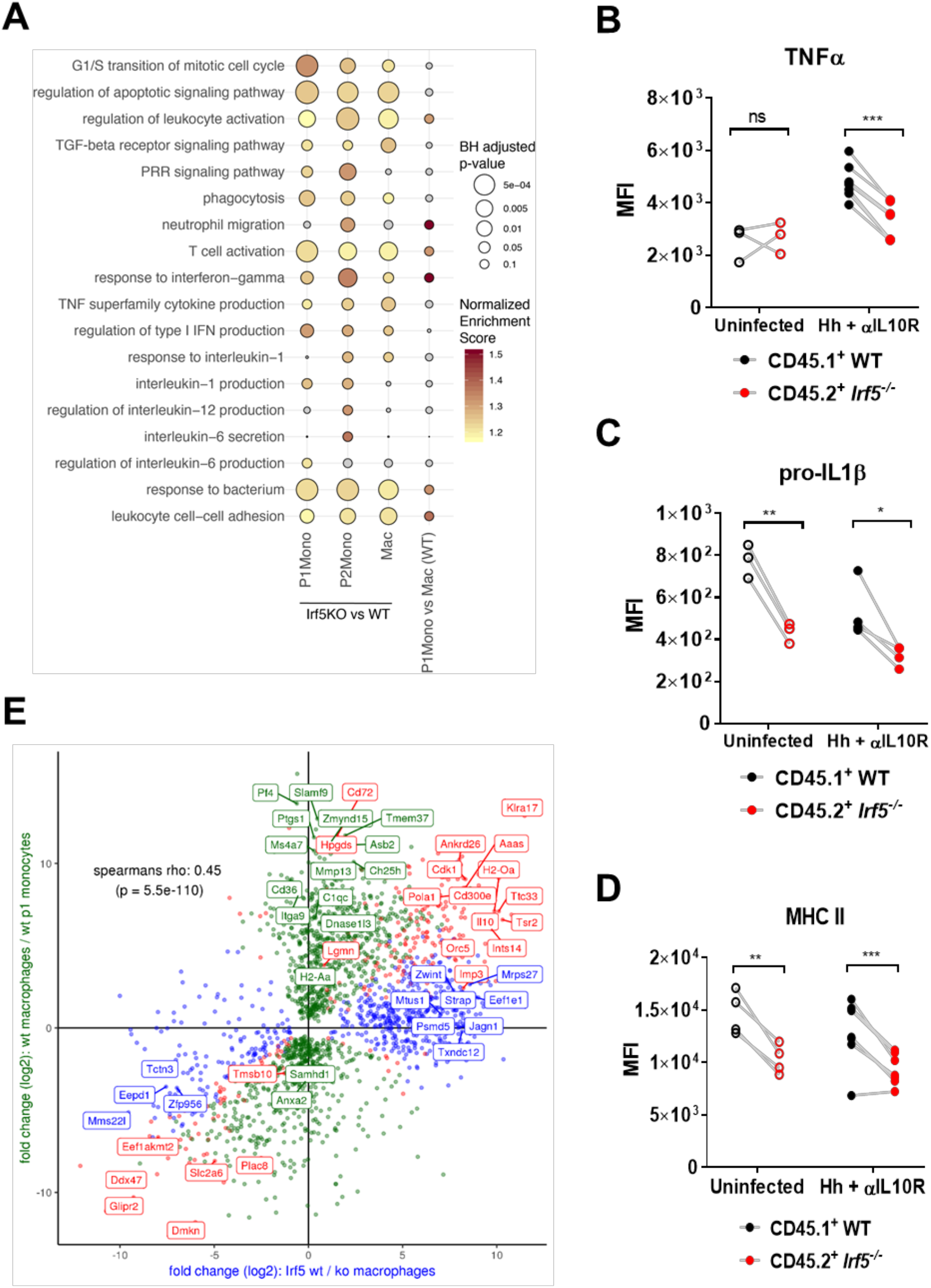
IRF5 endorses an inflammatory MNP phenotype. WT and *Irf5*^−/−^ P1 monocytes, P2 monocytes and macrophages were sorted from MBMC at d21 Hh + αIL10R. **A)** Selected GO Biological process categories that showed a significant enrichment (coloured dots, GSEA analysis, BH adjusted p < 0.1) in at least one of the three Irf5 KO vs WT small bulk RNA-seq comparisons. Intracellular or extracellular flow cytometry labelling was used to quantify the expression of **B)** TNFα, **C)** pro-IL1β, **D)** MHC II on WT vs *Irf5*^−/−^ macrophages in MBMC uninfected (n=3), and at d21 Hh + αIL10R colitis (n=4). **E)** Comparison of genes differentially expressed in WT vs *Irf5 ^−/ −^* macrophages with those differentially expressed between WT P1 monocytes and WT macrophages. Genes are coloured to indicate significant differential expression in both comparisons (red dots), in only WT vs *Irf5*^−/−^ macrophages (blue dots) or in only WT P1 monocytes vs WT macrophages (blue dots) (DESeq2 analyses, BH adjusted p < 0.05, see Supplementary Figure S4b). **A, B, C, D, E:** one experiment. **B, C, D)** Two-Way ANOVA with Sidak Correction Data presented are mean ± SEM, ns = not significant, * p ≤ 0.05, ** p ≤ 0.01, *** p ≤ 0.001

Next, we investigated the idea that the pro-inflammatory effect of Irf5 in the inflamed cLP might be explained by a deficiency in cellular differentiation. Assessment of the enrichment of the IRF5-regulated pathways amongst genes differentially expressed between the WT P1 monocytes and macrophages indicated that several of these pathways were also associated with monocyte differentiation (Fig. 4A, right column). We therefore performed a global comparison of genes regulated by IRF5 in macrophages with those that were associated with differentiation from P1 monocytes to macrophages in the inflamed cLP. Overall, this analysis revealed a significant positive correlation between genes up-regulated in macrophages and those positively regulated by IRF5 in macrophages (Spearman’s’ rho: 0.45, p = 5.5 × 10^−110^) (Fig. 4E and Supplementary Fig. S4B). Close examination of the scatter plot, however, also revealed a large number of genes that were up-regulated in macrophages independent of the presence of IRF5 including *C1qc*, *Ptgs1* (Cox1) and *Mmp13* (green dots, Fig. 4E). In summary, these data support a cell-intrinsic role for Irf5 in regulating myeloid cell differentiation and phenotype during intestinal inflammation. Through these roles, IRF5 controls the expression of a broad range of mediators of intestinal inflammation in populations of intestinal MNPs, with macrophages displaying the largest number of genes affected by IRF5 deficiency.

### Heterogeneity of MNP populations in inflamed intestine

At the population level, our data suggested roles for IRF5 in supporting the generation of intestinal macrophages and promoting pro-inflammatory gene expression (Fig. 3, Fig. 4, Supplementary Fig. S4C). To investigate the molecular basis of these phenomena in more detail, populations of CX_3_CR1^+^ WT and *Irf5*^−/−^ MNPs were isolated from the inflamed cLP of three Hh + αIL10R MBMCs and subjected to droplet-based single-cell RNA Sequencing (scRNA-seq) (Supplementary Fig. 5A). Following removal of low-quality cells and down-sampling to equalise cell numbers (n=553 cells retained per genotype) we used graph-based clustering (as implemented in the Seurat package, see methods) to identify eight distinct subpopulations (Fig. 5A). Examination of the top cluster markers genes (Supplementary Fig. S5B) revealed the existence of two groups of monocytes, two clusters of macrophages and four clusters of dendritic cells (Fig. 5A).

**Figure 5:**
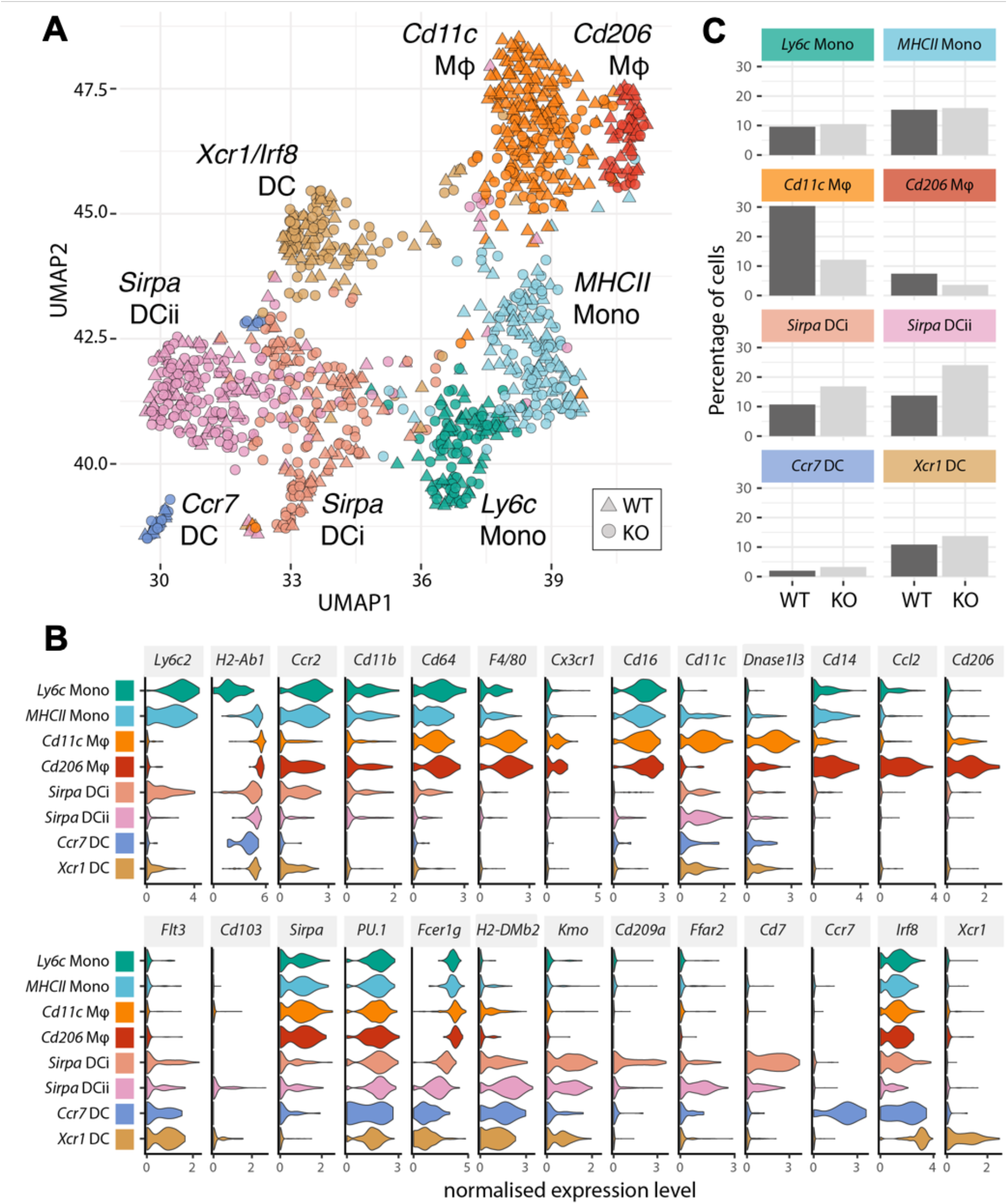
Heterogeneity of WT and *Irf5*^−/−^ CX_3_CR1^+^ MNPs in the cLP defined by massively parallel single cell RNA sequencing. WT and *Irf5*^−/−^ CD45^+^CD11b^+^SiglecF^−^Ly6G^−^CX_3_CR1^+^ cells were sorted from the colons of MBMC and subjected to droplet-based single cell transcriptomic analysis. **A)** Graph based clustering^76^ of equal numbers of WT and *Irf5*^−/−^ cells (n=1106 total) identified four clusters of MNPs and four clusters of dendritic cells. **B)** The violin plots show the expression levels (x axes) of selected known cLP MNP and DC sub-population markers in each of the identified clusters (y axes). **C)** The bar plots show the percentages of WT and *Irf5*^−/−^ cells that were found in each cluster.

The monocyte clusters, as identified by high expression of *Ccr2* and an absence of Mϕ and DC markers comprised of a set of *Ly6c2*-high immature monocytes (“*Ly6c* Mono”) and a group of mature monocytes (“*MHCII* Mono”) that expressed MHCII genes such as *H2-Ab1* and the pro-inflammatory cytokine *Il1b* (Fig. 5B). The two macrophage clusters were clearly demarcated by expression of *Adgre1* (*F4/80)*, *Cd81* and *Cx3cr1*. The largest cluster of *Cd11c* Mϕ was characterised by the high expression level of *Itgax* (*Cd11c*) and known cLP macrophage markers, such as MHC glycoproteins (*H2-M2*), complement molecules (*C1qa*,*b*,*c)*, tetraspanins (*Cd63*, *Cd72, Cd81*), oxidative stress response (*Hebp1*) and anti-microbial molecules (*Lyz2*, *Acp5, Dnase1l3*) (Fig. 5B and Supplementary Fig. S5B). The second cluster of *Cd206* Mϕ lacked *Itgax* expression but was defined by high expression of *Mrc1* (*Cd206*), chemokines (e.g. *Ccl2, Ccl3*, *Ccl4*, *Ccl7, Ccl12, Cxcl2)*, scavenger, phagocytic and immunoactivating receptors (*Cd36, Fcgr4, Clec4b1*) and anti-viral molecules (*Ch25h, Gbp2b*) and *Il-10* (Fig. 5B and Supplementary Fig. S5B). No *Timd4* (Tim-4) expression was detected in either of the macrophage clusters (Supplementary Fig. S5B), indicating that, as expected, the monocyte-independent resident macrophage population was not represented amongst the donor-derived cells^18^.

When scRNA-seq data were compared to the small bulk gene expression data, a good correspondence between the genes expressed in the *Ly6c* and *MHCII* monocyte clusters and P1 and P2 samples respectively was observed (Supplementary Fig S5C). Both the *Cd11c* and *Cd206* macrophage clusters showed similarities to the small-bulk macrophage sample (Supplementary Fig S5C).

The remaining four clusters of cells lacked *Cd64* expression and showed expression of established dendritic cell markers such as *Flt3*, *Cd11c*, and the *Ciita*-dependent DC-specific MHCII genes *H2-DMb2* and *H2-Oa*^42^. “*Sirpa* DC i” and “*Sirpa* DC ii” clusters displayed a DC2-like profile being marked by expression of *Sirpa*, *Kmo, Cd209a* and Cd7 (Fig. 5A **&** Supplementary Fig S5B). The cells in these cluster also strongly expressed *PU.1*, but were low for *Flt3*, suggesting that they may be monocyte-derived dendritic cells (moDC) ^43, 44^ (Fig 5). The remaining two DC clusters comprised of a set of *Xcr1*^high^*Irf8*^high^*Sirpa*^low^ cells (“*Xcr1* DC”) that are likely to correspond to conventional cDC1 cells and a small group of *Ccr7* positive (“*Ccr7* DC”) cells likely to represent migratory DCs.

When clusters were split by genotype, it was found that monocyte clusters had similar numbers of WT and *Irf5*^−/−^ cells, whereas macrophage clusters had higher numbers of WT cells (Fig. 5C) and the *Sirpa* DC i and ii clusters contained higher numbers of *Irf5*^−/−^ cells (Fig. 5C), consistent with our earlier observation (Fig 3).

### Generation of CD11c^+^ and CD206^+^ macrophage populations in the cLP

In order to investigate the shape of the monocyte-macrophage differentiation waterfall in the cLP we used the Slingshot pseudo-time algorithm^45^. In line with current understanding, this analysis revealed a continuum of differentiation from naïve *Ly6c* monocytes to *MHCII* monocytes to *Cd11c* macrophages to possibly *Cd206* macrophages (Fig. 6A). This was supported by pseudo-time calculated trajectory of expression of the waterfall characteristic markers, such as *H2-Aa* and *F4/80* (upregulated towards terminally differentiated mature macrophage state) and *Ly6c*2 (downregulated) (Fig. 6B). However, when we considered all genes that were differentially expressed along the trajectory, an apparent switch in gene expression between MHCII^+^ monocytes and macrophages was revealed, suggesting that the differentiation of monocytes into macrophages is likely to be a regulated cell fate choice rather than a gradual and inevitable transition (Supplementary Fig. S6A).

**Figure 6:**
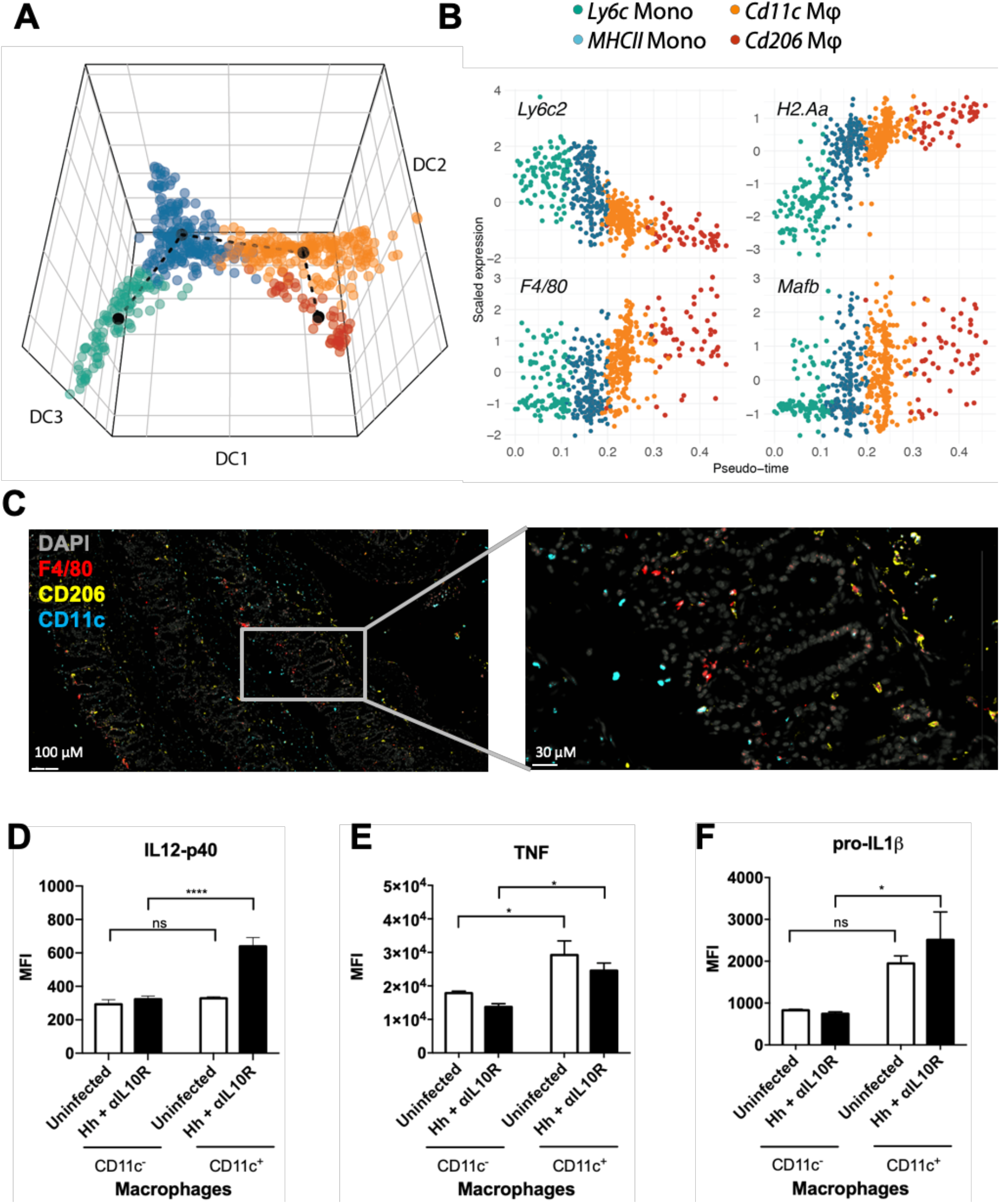
Monocyte differentiation in the cLP gives rise to *Cd11c* and *Cd206* macrophages. **A)** Embedding of the MNP cells in the first three dimensions of a diffusion map (DC1-3). The dashed line shows the minimum spanning tree computed by the pseudo-time algorithm Slingshot^45^, which identified a single trajectory from *Ly6c* monocytes (which were specified as the starting point) to *MHC II* monocytes, through to *Cd11c* macrophages and then *Cd206* macrophages. **B)** The scatter plots show the scaled expression of *H2-Aa*, *Ly6c2*, *Adgre1*, and *Mafb* in MNP cells ordered by their position in the identified trajectory (pseudo-time). **C)** Immunohistofluorescence to identify localisation of F4/80^+^ (Red), CD11c^+^ (Blue), and CD206^+^ (yellow) cells in the colons of steady state mice. Cell nuclei were labelled with Hoescht 33342 (Grey). **D-F)** Comparison of inflammatory cytokine expression in CD11c^+^ vs CD11c^−^ cLP macrophages assessed by intracellular flow cytometry, **D)** IL12p40, E) TNF, F) pro-IL1β in uninfected MBMC (n=3) and d21 Hh + αIL10R (n=4). **C)** Sections from 4 steady state mice were cut and labelled. A minimum of 5 sections per mouse were evaluated. **D, E, F)** One experiment. Two-Way ANOVA with Tukey Correction. Data presented are mean ± SEM, ns = not significant, * p ≤ 0.05, **** p < 0.0001

Although the minimum spanning tree inferred by Slingshot suggested that differentiation of *Cd11c* Mϕs may precede the emergence of *Cd206* Mϕs, other possibilities, such as an independent emergence of *Cd206* Mϕs cannot be excluded (Fig. 6A). In comparison to *Cd11c* Mϕs, *Cd206* Mϕs expressed higher levels of the transcription factor, MafB (Fig 6B), that has been shown to support macrophage terminal differentiation^46^. Confocal microscopy analysis of the CD11c^+^F4/80^+^ and CD206^+^F4/80^+^ macrophage location in the colon revealed that the macrophages occupy different geographical niches (Fig 6C). CD206^+^F4/80^+^ macrophages were found mainly at the base of crypts (Fig 6C). Macrophages at the base of crypts are believed to be anergic, IL10-expressing cells that sense luminal antigen via sampling between intestinal epithelial cells^47, 48^. In contrast, CD11c^+^F4/80^+^ macrophages were found throughout the Lamina Propria, including the tips of the villi. Thus, CD11c^+^ macrophage may represent a primed macrophage phenotype, ready to respond to microbial encroachment^49^. In fact, CD11c^+^F4/80^+^ macrophages are critical effector cells in the development of experimental colitis via the production of IL-23^9^. Indeed, during Hh + αIL10R-induced colitis CD11c^+^F4/80^+^ macrophages produced higher levels of IL-12p40, a subunit of IL-23, than CD11c^−^ macrophages (Fig. 6D). Production of other inflammatory cytokines, such as TNF and IL-1β was also elevated in CD11c^+^F4/80^+^ macrophages (Fig. 6E, F). Moreover, during inflammation the balance of macrophage populations was shifted towards the predominance of CD11c^+^CD206^−^ macrophages (Supplementary Fig. S6B).

### IRF5 marks and controls the pathogenic CD11c^+^ macrophage population

WT and *Irf5*^−/−^ cells were distributed differently along the monocyte to macrophage differentiation trajectory with *Irf5*^−/−^ cells tending to accumulate significantly earlier in pseudo-time than their WT counterparts (Wilcoxon test, p < 3.48 × 10^−8^) and to give rise to proportionally fewer *Cd11c* Mϕs (Fig. 7A). Flow cytometry analysis of macrophage populations confirmed a significant loss of *Irf5*^−/−^ CD11c^+^F4/80^+^ macrophages in MBMCs during during Hh + aIL10R-induced colitis, with no substantial effect on generation of CD206^+^F4/80^+^ macrophages (Fig. 7B). Similarly, *Irf5*^−/−^ CD11c^+^F4/80^+^ but not *Irf5*^−/−^ CD206^+^F4/80^+^ macrophages were reduced in MBMCs in the absence of inflammation (Supplementary Fig. S7A). Taking into account equal frequencies of WT and *Irf5*^−/−^ Ly6C^hi^MHCII^−^ (P1) and Ly6C^hi^MHCII^+^ (P2) monocytes in the MBMCs (Fig. 3A,B) and unaltered apoptosis rates of macrophages (Fig. 1F), these data suggest that IRF5 may control cell fate choice of Ly6C^hi^MHCII^+^ monocytes promoting their acquisition of CD11c expression and transition to pro-inflammatory CD11c^+^F4/80^+^ macrophage phenotype. In fact, we found the majority of IRF5^+^ Ly6C^hi^MHCII^+^ monocytes and macrophages to be CD11c^+^ cells (Supplementary Fig. S7B, C) and the majority of CD11c^+^ P2 monocytes and macrophages to be IRF5^+^ cells (Supplementary Fig. S7D, E). In addition, the expression of IRF5 protein was higher in CD11c^+^ macrophages as well as CD11c^+^ Ly6C^hi^MHCII^+^ monocytes compared to their CD11c^−^ counterparts, both at steady state and during Hh + αIL10R colitis (Fig. 7C, Supplementary Fig. S7F).

**Figure 7:**
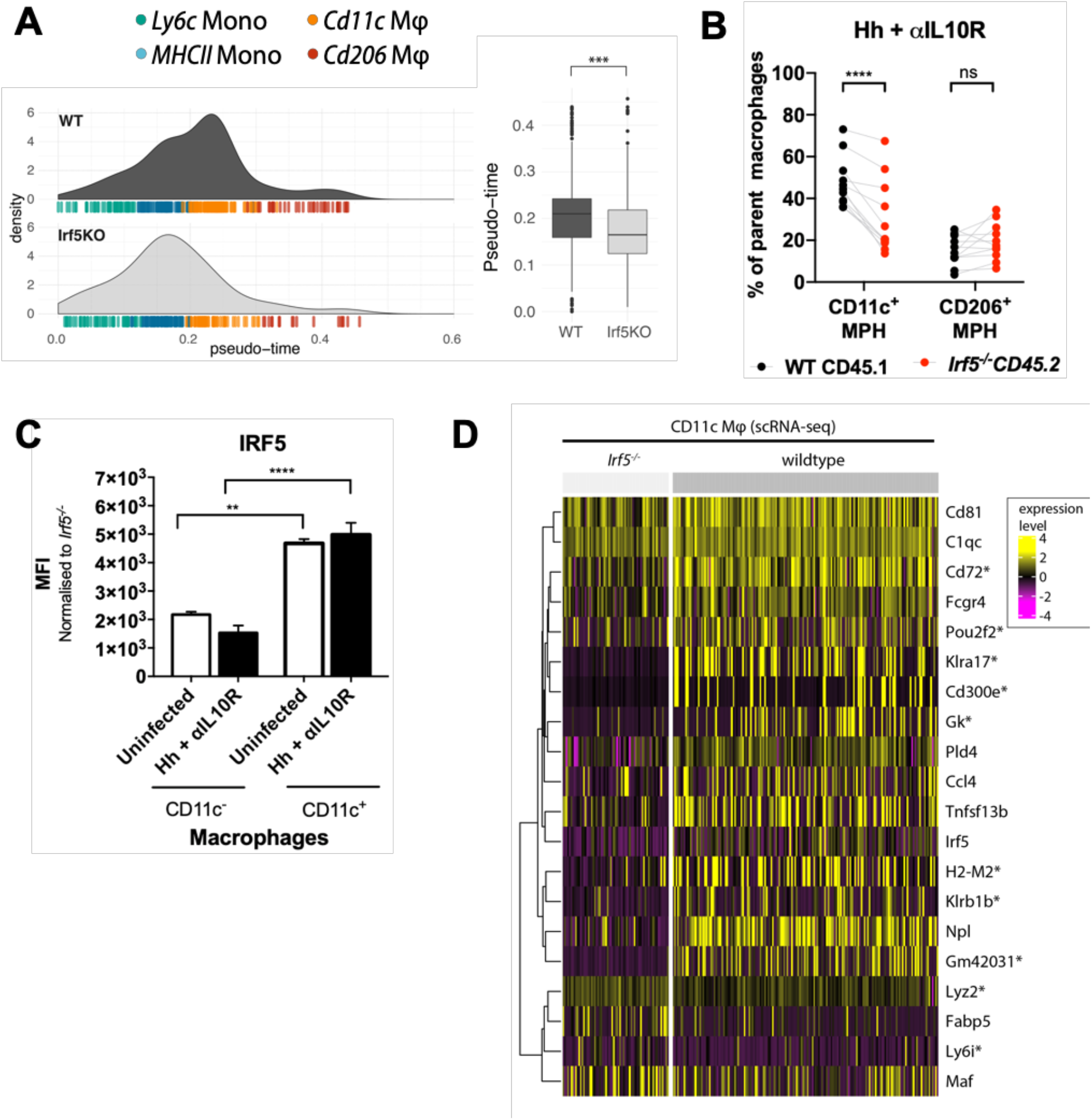
IRF5 specifies a pathogenic CD11c^+^ macrophage phenotype during intestinal inflammation. **A)** The density plots (left) and boxplots (right) summarise the distribution of individual WT and Irf5^−/−^ MNP from the inflamed cLP of MBMCs along the identified differentiation trajectory (see Figure 6a). The WT cells had significantly higher pseudo-time scores (two-tailed Wilcoxon test, p=3.48 × 10^−8^). **B)** WT CD11c^+^ macrophages, but not CD206^+^ macrophages make up are greater frequency of parent macrophages in MBMC (n=12). IRF5 expression in CD11c^+^ vs CD11c^−^ **C)** P2 monocytes, **D)** macrophages in MBMC assessed by intracellular flow cytometry. **E)** The heatmap shows expression of selected genes in WT and *Irf5^−/−^ Cd11c* macrophages from the inflamed cLP of the MBMCs (see Figure 5). All of the genes shown were found to be significantly differentially expressed between the WT and *Irf5*^−/−^ cells of this cluster (Wilcoxon tests, BH adjusted p < 0.05). *’s denote significant differential expression between the genotypes in the macrophage small-bulk RNA-seq data. Frequency of CD11c^+^ within IRF5^−^ of IRF5^+^ **F)** P2 monocytes, and **G)** macrophages in the cLP. **C, D, F, G:** one experiment, uninfected n=3, Hh + αIL10R n=4. **B,F,G)** Two-Way ANOVA with Sidak Correction. **C,D)** Two-Way ANOVA with Tukey Correction. Data presented are mean ± SEM, ns = not significant, * p ≤ 0.05, ** p ≤ 0.01, *** p ≤ 0.001, **** p < 0.0001

Thus, IRF5 imposed control of intestinal inflammation (Fig 2) is likely to be due to its function in CD11c^+^ macrophages. In these cells IRF5 positively regulated genes that defined cLP macrophage phenotypes, such as MHC molecules (*H2-M2*), tetraspanins (*Cd72, Cd81*), complement molecules (*C1q*), chemokines (*Ccl4*), metalloproteinase (*Acp5*) and phagocytic and immunoactivating receptors (*Fcgr4, Fcer1g, Cd300e*). A number of killer cell lectin-like receptor family members (*Klrb1b, Klra2, Klra17*), not previously associated with macrophage function were also affected by the lack of IRF5 in this compartment (Fig 7D and Supplementary Fig S5B).

Together, these data demonstrate that IRF5 marks, induces and maintains pathogenic CD11c^+^ intestinal macrophage population during Hh + αIL10R-induced colitis.

## Discussion

Using a model of mononuclear phagocyte development in the gut and a combination of mixed bone marrow chimera approaches and single cell analysis of gene expression, we identify a novel function for IRF5 in promoting the generation of macrophages in the cLP, marking a pathogenic CD11c^+^F4/80^+^ macrophage phenotype in inflammation, and controlling the immunopathology of Hh + αIL10R-induced colitis.

The TLR4/Myd88 signalling cascade is important for the establishment of intestinal homeostasis^40^ and MyD88 is required for the development of colitis in *Il10*^−/−^ deficient mice^50^. IRF5 is a known target of MyD88, and we found no evidence that the MyD88 signalling cascade was disrupted in the absence of IRF5 (Supplementary Fig. S3) in our experiments^51–53^. The fact that mice with both global or MNP-specific loss of IRF5 showed protection from Hh + αIL10R colitis (Fig. 2) thus strongly argues that initiation of intestinal colitis in this model is a consequence of TLR signalling via MyD88 in myeloid cells, and that IRF5 is essential for this process.

To assess the role of IRF5 in the inflamed intestine, we took advantage of MBMC. The presence of Irf5-deficient cells did not fully abolish inflammation in the Hh +aIL10R model, but rather revealed a cell-intrinsic role for IRF5 in the control of genes and biological pathways related to monocyte differentiation, leukocyte activation, response to bacterium, pattern recognition receptor signalling pathway, regulation of T-cell activation, (Fig. 4 **and** S4C). In particular, a major function of IRF5 identified in this study in the inflamed intestine is its ability to promote a pro-inflammatory macrophage state, which is positive for CD11c. Notably, inflammation induced an accumulation of CD11c^+^ macrophages in the cLP, which were also marked by high level of IRF5 (Supplementary Fig S6, S7). Irf5 deficiency ameliorated this induction (Fig 7). CD11c^+^ macrophages, which are highly bactericidal, produce high quantities of IL23 in the early stages of colitis and are essential for triggering intestinal immunopathology^9, 15^. They can be found throughout the cLP, and are efficient producers of inflammatory cytokines, such as TNF and IL-1β, that support pathogenic T cell responses in the intestine^33, 54, 55^ (Fig 6). Being intimately related to MHCII^+^ monocytes in the continuum of monocyte to macrophage differentiation, CD11c^+^ macrophages are likely to represent what was previously described as monocyte-macrophage intermediates and effector cells in multiple inflammatory disorders^56, 57, 58, 59^. With the recently established link between CD11c^+^ macrophages and IRF5 in the development of atherosclerotic lesions^60^, our data here support the notion that IRF5 may mark and maintain pathogenic CD11c^+^ macrophages in a variety of tissues and pathologies.

Another population of macrophages detected in our analyses is CD206-positive macrophages, which are located at the base of crypts (Fig 6), express MafB (Fig 6) and IL10 (Supplementary Fig S5). These macrophages are likely to represent fully differentiated tolerogenic macrophages involved in sensing luminal antigen via sampling between intestinal epithelial cells, and contributing to epithelial renewal via the regulation of *Lgr5^+^* stem cells^13, 61^. The *Cd206* macrophages appeared largely unaffected by IRF5 deficiency (Fig 7) and unlikely to be major contributors to the Hh-induced pathology^32^. The relationship between *Cd11c* and *Cd206* macrophage populations remains elusive. While pseudo-time analysis of our scRNA-seq data indicated that *Cd206* macrophage may represent a post-*Cd11c* stage of macrophage differentiation (Fig 5), the distinct distribution of the two macrophage populations in the cLP (Fig 6) and their unequal dependence on IRF5 (Fig 7) suggest that they may emerge independently in specific environmental niches. Indeed, two recent publications have described the predominance of two subpopulations of macrophages identified by scRNAseq in multiple tissues with distinct localisations^62, 63^. Monocytes were shown to develop into two distinct lineages of macrophage (Lyve1^lo^ MHCII^hi^ and Lyve1^hi^ MHCII^lo^), likely dependent on specific environmental cues ^62^. Interestingly, Lyve1^lo^ MHCII^hi^ macrophages also highly expressed markers in common with our *Cd11c* macrophages, including Itgax, *Cd9*, *Cd52*, *Cd72* and *Cd74*, whilst Lyve1^hi^ MHCII^−^ macrophages expressed *Mafb* and *Mrc1* (*Cd206*) in common with our *Cd206* macrophages ^62^.

Throughout the MNP compartment of the inflamed intestine IRF5 broadly modulated biological pathways related to leukocyte activation, response to bacterium, pattern recognition receptor signalling pathway and regulation of T-cell activation (Fig. 4). Thus, our data suggest that, subsequent to initiation of inflammation, TLR signalling via MyD88 promotes colitis in part by dictating IRF5-dependent pro-inflammatory transcriptional programs throughout the MNP. Indeed, mice deficient in IRF5 globally or in the MNP compartment only were protected from Hh + αIL10R colitis (Fig. 2).

Migration of leukocytes into sites of inflammation is a critical step in disease pathogenesis. The *Cd206* macrophages in the cLP readily produce CCL2 (Fig. 5), a critical chemokine for accumulation of monocytes in the cLP^20^. WT monocytes accumulated more efficiently in the cLP and migrate more readily towards CCL2 in an *in vitro* migration assay, than *Irf5*^−/−^ monocytes (Supplementary Fig. S3). These data support an intrinsic role for IRF5 in guiding the recruitment of monocytes to the primary site of inflammation^36^ and highlight another mechanism by which IRF5 could modulate inflammation *i.e*. via controlling a pathogenic positive-feedback loop of inflammatory monocyte recruitment.

Finally, our *in vivo* data (Fig. 3, 4, 5) and *in vitro* bone marrow culture (Fig. 3) suggest that in an inflammatory environment, IRF5 specifically promotes key aspects of macrophage transcription whilst repressing key DC genes. Given the observed changes in expression of the histone deacetylases *Hdac2* and *Hdac9*, it seems likely that these actions involve modulation of the epigenetic state of these cells. This process may be due to loss of competition for IRF binding sites, and engagement of an IRF4-dependent differentiation program^64, 65^. IRF4 and IRF5 were shown to compete for binding to Myeloid Differentiation primary response 88 (MyD88) and activation following TLR4 ligation^66^. IRF4 is a key regulator of intestinal CD11b^+^ DC subsets and a critical transcription factor in the DC fate of monocytes in *in vitro* bone marrow cultures^65, 67^. Thus, in the absence of IRF5, IRF4 may be able to dominate the fate choice of monocytes, explaining the increased predisposition to DC fate in *Irf5*^−/−^.

Although intestinal DCs are believed to be largely derived of FLT3L-dependent progenitors^13^, several studies have provided evidence that *Sirpa* CD11b^+^ DCs are replenished by monocytes in the inflamed cLP^16, 17^. Our attempts to adoptively transfer WT and *Irf5*^−/−^ monocytes in non-perturbed intestinal environment had a limited success, as only a few transferred cells were detected in the colon, while perturbations that improved on transfer efficiency (e.g. transfer into CCR2-deficient mice) had a strong effect on the system and were thus deemed uninformative. Other niche-depleting strategies may help to clarify this issue.

In summary, the data presented here reveals that IRF5 is essential for the initiation of intestinal inflammation and acts as a cell-intrinsic master regulator of pathogenic myeloid cell differentiation and phenotype during intestinal inflammation.

## Materials and Methods

### Study Design

The purpose of this study was to understand the intrinsic role of IRF5 in directing macrophage polarisation and intestinal inflammation. We used flow cytometry, bulk- and single cell-RNA-sequencing, and immunofluorescence labelling of intestinal tissue sections to analyse the leukocyte milieu in the colons of wild type or *Irf5*^−/−^ or mixed bone marrow chimeric mice. Mice were aged between 8-16 weeks at the commencement of experiments. Experimental sample sizes were not predetermined. *Helicobacter hepaticus* infections were ended upon mouse sacrifice at d21 post-infection. Bone marrow-derived macrophages were cultured for 8 days and harvested post stimulation on d9. Experimenters. In general, experiments were performed twice unless indicated otherwise. Data were not excluded from analysis except for qc failures in RNAseq analysis detailed below. Histopathology assessment was conducted in a blinded manner independently by two researchers. Experimenters were not blinded to intervention groups for flow cytometry analysis.

### Animals

Mice were bred and maintained under SPF conditions in accredited animal facilities at the University of Oxford. All procedures were conducted according to the Operations of Animals in Scientific Procedures Act (ASPA) of 1986 and approved by the Kennedy Institute of Rheumatology Ethics Committee. Animals were housed in individually ventilated cages at a constant temperature with food and water *ad libitum*. Mice were free of known intestinal pathogens and negative for *Helicobacter* species. C57Bl/6 and Irf5^tm1/J^ (*Irf5*^−/−^) mice were bred in house. C57Bl/6 SJL CD45.1^+^ WT mice were purchased from the University of Oxford BMS.) Cx3Cr1^tm1.1(cre)Jung/J^, (CX_3_CR1^IRF5+^, cWT) were acquired from JAX (Jackson Laboratories, UK, JAX stock #025524)) and crossed with C57BL/6-Irf5^tm1Ppr^/J (Irf5^fl/fl^) to generate conditional *Irf5*^−/−^ mice (cKO, CX_3_CR1^IRF5−^). CX_3_CR1^GFP/GFP^ CD45.1^+^ mice (B6.129P(Cg)-Ptprc^a^ Cx3cr1^tm1Litt/LittJ^) were maintained in house. B6.SJL-*Ptprc*^a^ (CD45.1^+^ WT mice) were crossed with B6.129P(Cg)-Ptprc^a^ Cx3cr1^tm1Litt/LittJ^ to generate CD45.1^+^ CX_3_CR1^GFP/+^, and B6.129P-Cx3cr1tm1Litt/J were crossed with Irf5^−/−^ to generate CD45.2^+^ CX_3_CR1^GFP/+^ Irf5^−/−^ mice.

### Generation of Bone Marrow Chimaeras

≥ 20 g male C57BL/6 SJL (CD45.1^+^ WT) mice were selected as bone marrow recipients. 2 days before irradiation, mice were fed antibiotic-treated water (Co-trimoxazole, Aspen Pharma), or Baytril, Bayer) which was maintained until 2 weeks post-irradiation (PI). Recipients were lethally irradiated by exposure to 11 Gy split into two equal doses 6 hrs apart in an X-ray irradiator (Gulmay). Within 24 hrs post-irradiation, 4 x10^6^ freshly isolated bone marrow cells at a 1:1 ratio (CD45.1^+^ WT and CD45.2^+^ *Irf5*^−/−^, or CD45.1^+^ CX_3_CR1^GFP/+^ and CD45.2^+^ CX_3_CR1^GFP/+^ Irf5^−/−^) were injected i.v. into the tail vein of CD45.1^+^ WT hosts.

6 weeks post-irradiation, reconstitution was assessed by collecting blood leukocytes by tail vein bleed.

### Helicobacter hepaticus culture

*Helicobacter hepaticus* (Hh) NCI-Frederick isolate 1A (Hh1a; strain 51449; American Type Culture Collection) was grown at 37°C on blood agar plates containing Campylobacter-selective supplement (skirrow), (Oxoid) under microaerophilic conditions (10% CO_2_, 10% H_2_, balance N_2_) in a vented CampyPak jar (Oxoid).

After 2 days of blood agar plates, cultures were harvested using cotton swabs and transferred to liquid culture (Tryptan Soya Broth (Sigma Aldrich) dissolved in 1000 mL MilliQ H_2_O, and autoclaved) containing 10% FCS + 4 mL Campylobacter-selective supplement (skirrow) (Oxoid). Hh culture was inoculated at 0.05 OD_600_ in a 500 mL vented Erlenmeyer flasks (Corning). Liquid culture was maintained at 37°C shaking at 100 rpm under microaerophilic conditions, and split every 24 hours to 0.05 OD_600_ to maintain stable growth.

Hh viability was assessed by labelling with a fluorescent live/dead assay kit (Life Technologies) according to the manufacturer’s protocol using a CKX41 fluorescence microscope (Olympus).

### *Helicobacter hepaticus* + αIL10R model of colitis

Mice were infected with 1×10^8^ colony forming units (cfu) Hh in 200 μL sterile PBS on days 0 and 1 by oral gavage with a 22G curved, blunted needle (Popper & Sons). Mice were injected intraperitoneally once weekly starting on day 0 with 1 mg anti-IL10R blocking antibody (clone 1B1.2) in a volume of 200 μL. Infected mice were monitored weekly for colitis symptoms. Mice were culled by Schedule 1 method three weeks after day of infection, and organs were harvested for analysis.

### Assessment of bacterial colonisation

Caecal contents were collected after mice were sacrificed. DNA was isolated from faeces using Stool DNA extraction kit (Qiagen) as per manufacturer’s instructions. qPCR was performed with primers against the *Hh cdtB* gene using a Viia7 Real-Time PCR system (Applied Biosystems) as described by Maloy *et al.*, (2003) ^68^. In order to construct standard curves, DNA was extracted from Hh cultures using a DNeasy Kit (Qiagen). Primer sequences: *cdt*B Reverse - *TCG TCC AAA ATG CAC AGG TG, cdt*B Forward - *CCG CAA ATT GCA GCA ATA CTT*, *cdt*B Probe - *AAT ATA CGC GCA CAC CTC TCA TCT GAC CAT*.

### qPCR

For each reaction, 10 ng cDNA was added to 2.5 μL 2X qPCR FAST mastermix (Primerdesign). 0.1 μL primer + Taqman probe mix (Table.M2) was added to each sample, and the reaction volume was topped up to 5 μL with RNase/DNase-free H_2_O (Promega). Thermal cycling was carried out using a Viia7 real time PCR system (Applied Biosystems). Thermal cycling: 1×120 s95°C, 40x(5 s 95°C 20 s 60°C).

### Histopathological assessment

Post-sacrifice, 0.5 cm pieces of caecum, and proximal, mid and distal colon were fixed in PBS + 4% paraformaldehyde (Sigma Aldrich). Fixed tissue was embedded in paraffin blocks, and sectioned using a microtome and stained with Haematoxylin and Eosin (H&E) by the Kennedy Institute of Rheumatology Histology Facility (Kennedy Institute of Rheumatology, University of Oxford).

Sections were scored in a blinded manner by two researchers according to Izcue *et al*., (2008) ^69^.

### Immunofluorescence labelling of colons

After whole colon excision and longitudinal slicing, colon tissue was washed in PBS, rolled into Swiss rolls, embedded in Optimal Cutting Temperature (OCT) medium (Tissue-Tek), before freezing on dry ice with 2-methylbutane, and stored at −80°C (Bialkowska et al. 2016). 5 μM cryosections were air dried and rehydrated with PBS. The sections were fixed in a 1:1 mixture of methanol:acetone (Merck) and blocked with PBS containing 5% goat serum and 5% mouse serum (blocking solution). Sections were then blocked with biotin/avidin (Invitrogen). Sections were then labelled with primary antibodies (α-F4/80 (recombinant CI:A3-1, Enzo), α-CD11c-biotin (N418, Biolegend), α-CD206 (MR5D3, Bio-rad)) performed in blocking solution overnight at 4°C. Secondary antibody labelling was performed with goat α-rat-Alexa Fluor 488, goat α-rabbit-Alexa Fluor 555 and streptavidin-Alexa Fluor 647 (all Thermo Fisher). The sections were stained at RT in the dark for 30 min. Sections were then stained with Hoechst 33342 (Thermo Fisher) for 10min. After PBS washes, the tissue was mounted using 5% N-propyl gallate (Merck) in glycerol and imaged using a Zeiss Axio Scope A1 (Zeiss). The images were analysed using IMARIS.

### Isolation of lamina propria leukocytes

Colons and/or caeca were harvested from mice, washed in PBS/BSA and content flushed with forceps. Intestines were then opened longitudinally and washed once more before blotting to remove mucus. Gut tissue was then cut into 1 cm long pieces and placed in 50 mL centrifuge tube (Greiner) in ice cold PBS + 0.1% BSA. Colons were incubated 2 times at 200 rpm in 40 mL HBSS + 0.1% BSA + 1% Penicillin-Streptomycin (PS, Lonza) + 5mM EDTA (Sigma-Aldrich) at 37 °C for 10 min before the supernatant was aspirated. Tissue was placed in 40 mL PBS + 0.1% BSA + 1% PS for 5 min. Intestines were then incubated with 20 mL RPMI + 10% FCS +1% PS + 2.5 U/mL Collagenase VIII (Sigma-Aldrich) + 2 U/mL DNAse I (Roche), shaking at 200 rpm for 45 mins - 1 hour at 37 °C. Supernatant was filtered through a 70 μm cell strainer to which 30 mL of ice cold PBS + 0.1% BSA + 1% PS + 5 mM EDTA was added to ablate collagenase/DNase activity. Cells were washed in 30 mL PBS/BSA before filtering once more through a 40 μm cell strainer. The cells were then pelleted by centrifugation at 400 rcf for 10 minutes at 4 °C.

Colonic lamina propria leukocytes (cLPLs) were isolated by resuspending cells in 4 mL P80 (80% P100 (9:1 percoll:10X PBS) + 20% RPMI) percoll in a 15 mL centrifuge tube (Greiner) before overlaying 4 mL P40 (40% P100 + 60% 1X PBS) layer. Cells were spun at 3000 rpm for 20 min at room temperature, slow acceleration, no brake. Mucus and cellular debris were aspirated from the surface of the P40 layer with a Pasteur pipette and pipetted into 40 mL of ice cold PBS + 0.1% BSA. Cells were pelleted by centrifugation at 400 rcf for 10 min and resuspended in 1 mL RPMI + 10% FCS + 1% PS before counting.

### Isolation of blood leukocytes

Blood was harvested by either tail vein bleed or cardiac puncture. Mice were culled by Schedule 1 method in accordance with the project licence. Prior to cardiac puncture a 1 mL syringe was coated with PBS + 2 mM EDTA. Cardiac puncture was performed with a 27G needle.

Tail vein bleeds were performed using a #24 blade scalpel (Swann-Morton Ltd.)

Collected blood was placed in 1 mL of sterile 2 mM EDTA/PBS solution in a 15 mL centrifuge tube (Greiner). Tubes were topped up with ice-cold PBS + 0.1% BSA. Cells were pelleted by centrifugation at 400 rcf for 10 mins at 4 °C and the supernatant discarded. Erythrocytes were then lysed using 10-20X the blood sample volume of ACK lysis buffer (Gibco) for 3 mins. Tubes were then topped up to 15 mL with ice cold PBS + 0.1% BSA to quench the lysis buffer. Cells were washed in PBS/BSA, resuspended in PBS + 0.1% BSA to the desired cell concentration, and stored at 4 °C until required.

### Splenocyte isolation

Spleens were harvested from mice and stored in RPMI + 2% FCS at 4 °C until required. Spleens were then mashed through a 70 μm strainer, and washed through with 10 mL ice cold PBS + 1 mM EDTA. Cells were pelleted by centrifugation for 10 mins at 400 rcf at 4 °C and the supernatant was discarded. Erythrocytes were lysed as described above. Cells were resuspended in ice cold PBS + 0.1% BSA until required.

### Bone marrow isolation

Whole hind legs of mice were harvested and stored at 4 °C until processing. Processing was carried out in a class II lamina flow hood. Femurs and tibia muscle tissue was removed using scissors, followed by desiccation in 70% ethanol solution for 3 min. Remaining muscle was cleaned from the bones using a lint-free tissue.

Scissors were used to cut the ends of bones. A 27G needle was inserted into the opening, and marrow was flushed into a 50 mL centrifuge tube using 10 mL of sterile, ice cold PBS. Cells were then filtered through a 70 μm cell strainer (Greiner). Red blood cells were lysed with ACK as described above. Cells were resuspended in PBS + 0.1% BSA at 4 °C until required.

### Bone Marrow Macrophage differentiation

Isolated (non-RBC-lysed) bone marrow cells were plated at a density of 5×10^5^ cells/mL in RPMI-1640 + 10% FCS + 100 U/mL penicillin/streptomycin + 1 mM β-mercaptoethanol (BM medium) with 20 ng/mL murine GM-CSF (Peprotech) and cultures in a humidified environment at 37°C and 5% CO_2_. On day 3 of culture, fresh BM media with 20 ng/mL murine GM-CSF was added to cultures. On day 6, half of the culture media was harvested and spun at 400 rcf for 5 min to pellet cells. The supernatant was then discarded, and the pellet resuspended in fresh BM medium with 20 ng/mL murine GM-CSF. On day 9, mature macrophages were harvested by incubation with PBS + 0.48 mM EDTA for 5 min at 37°C.

### Monocyte isolation

Splenocytes were prepared as described above. Monocytes were enriched by negative selection using an EasySep™ Mouse Monocyte Isolation Kit (StemCell) according to the manufacturer’s instructions. Cells were then resuspended in ice cold PBS + 0.1% BSA for counting and downstream processing.

### T cell restimulation

A minimum of 1 x10^6^ freshly isolated cLPLs were incubated in V-bottomed 96 well plates (Corning) in 250 μL Iscove’s Modified Dulbecco’s Medium (Gibco) supplemented with 10% FCS + 1% P/S + 0.1 μM phorbol 12-myristate 13-acetate (PMA, Sigma Aldrich) + 1μM Ionomycin (Sigma Alrich) + 10 μg/mL GolgiPlug (BD) for 4 hrs at 37 °C. Cells were washed 3 × in 150 μL FACS buffer before extracellular and intracellular staining.

### Flow cytometry

CompBeads (BD) were used to prepare single stained controls as per manufacturer’s instructions to set up fluorophore compensation. Labelled cells were acquired using either an LSR II (BD), or Fortessa X20 (BD) flow cytometer using FACSDiva (BD). Data were analysed using Flowjo (Treestar, Inc.) software.

### Extracellular labelling of cells

5×10^5^ − 2×10^6^ cells were plated on either V-bottomed or U-bottomed 96 well plates. The cells were washed twice with 150 μL FACS buffer (PBS + 0.1 % BSA + 1 mM EDTA + 0.01% Sodium Azide) at 400 rcf for 3 min 4°C. Cells were then Fc blocked for 10 min with αCD16/CD32 (BD) 1/100 in 20 μL FACS buffer at room temperature (RT) followed by washing once in 150 μL FACS buffer. Fixable Viability Dye eFluor®780 (ThermoFisher) and primary extracellular antibodies were added for 20 min at 4 °C in 20 μL FACS buffer in the dark. Labelled cells were then washed twice with 150 μL FACS buffer. Cells were then fixed for 30 mins in 50 μL Cytofix (BD), washed twice with 150 μL FACS buffer, and resuspended in 200 μL FACS buffer before acquisition.

### Intracellular labelling of cytokines

Surface markers were labelled as described above. For Cytokine labelling, cells were fixed in 50 μL Cytofix/Cytoperm (BD), washed twice in 150 μL Perm/Wash buffer (BD) at 600 rcf, 4 °C. Intracellular antibodies were incubated in 20 μL Perm/Wash for 20 mins at 4 °C in the dark, after which samples were washed twice in 150 μL Perm/Wash, and once in 150 μL FACS buffer before resuspension in 200 μL FACS buffer for acquisition.

### Intracellular labelling of transcription factors

Surface markers were labelled as described above. For nuclear staining, cells were fixed in 50 μL Fix/Perm (eBioscience) according to manufacturer’s instructions, washed twice in 150 μL Perm buffer (eBioscience) at 600 rcf, 4 °C. Intracellular antibodies were incubated in 20 μL Perm for 20 mins at 4 °C, protected from light. Next, samples were washed twice in 150 μL Perm, and once in 150 μL FACS buffer before resuspension in 200 μL FACS buffer for acquisition.

### Cell sorting

Cells were stained with extracellular cytokines as described above, except no fixation step was performed. Cells were then sorted using a FACSAria™ II (BD) at the Kennedy Institute of Rheumatology FACS facility.

### RNA sequencing of GM-BMDMs

Total RNA from 1×10^6^ cells was isolated using RNeasy Mini kit (Qiagen). 50 b.p. single end reads were generated and mapped using STAR with the options: “--runMode alignReads --outFilterMismatchNmax 2.” HTSeq-count was then used to count uniquely mapped reads over annotated genes with the options: “-a 255 -s no -m union -t CDS” and differential expression was tested with DESeq2, with the following cutoffs: fold change > 1.5 and an FDR > 0.05.

Transcripts per million (TPM) were calculated with Salmon (Patro et al., 2017) from unmapped reads using quasi-mapping for the mouse mm10 assembly. Published Immgen datasets were used to identify macrophage ^39^ and DC ^38^ signatures.

### Generation and analysis of “small bulk” RNA-sequencing data from MBMC

For “small bulk” RNA-sequencing 100 cell samples were sorted through a 100 μm diameter nozzle into 2 μL of lysis buffer and amplified cDNA prepared using the Smart-seq2 protocol ^70^. Libraries were prepared using Nextera XT kits (Illumina) and sequenced to a mean depth of 17M read pairs (Illumina HiSeq 4000). Sequence reads were aligned to the mouse genome with Hisat2 (version 2.1.0) using a “genome_trans” index built from the mm10 release of the mouse genome and Ensembl version 91 annotations (two-pass strategy to discover novel splice sites; with parameters: --dta and --score-min L,0.0,-0.2) ^71^. Mapped reads were counted using featureCounts (Subread version 1.6.3; Ensembl version 91 annotations; default parameters) ^72^. TPMs were estimated with salmon (version 0.11.3, with parameter “--gcBias”) using a quasi index built from the full set of Ensembl version 91 transcriptome annotations (parameters “--k=31 --keepDuplicates”)^73^. The median alignment rate was 83.8% (assessed with Picard tools v2.10.9, https://github.com/broadinstitute/picard).

For each of the GSEA analyses (Figure 4A) genes detected in a least one of the biological replicates (n=3) were pre-ranked by p-value for differential expression (Deseq2, paired design, local fit for dispersion, no independent filtering). Enrichment of GO biological processes (org.Mm.ed.db R package version 3.7.0; with ≥5 and ≤500 genes) was tested using fgsea^74^ (version 1.8.0). Changes in gene expression across the wildtype monocyte waterfall (Figure S4A) were identified using DESeq2 (likelihood ratio test, modelling animal and cell-type). Differential expression analyses (Figure 4E, Supp. Figure 4B and C) were performed using DESeq2 (paired tests, local dispersion fit).

### Generation and pre-processing of single-cell RNA-sequencing data

Single-cell RNA-sequencing libraries were generated using the 10x Genomics Single Cell 3’ Solution (version 2) kit and sequenced to an average depth of > 200k reads/cell (Illumina HiSeq 4000). Data analysis was performed using Python3 pipelines (https://github.com/sansomlab/tenx) written using CGAT-core^75^. Read mapping, quantitation and aggregation of sample count matrices was performed with the 10x Genomics Cell Ranger pipeline (version 2.1.1). For the “cellranger count” step, a custom reference was built using Ensembl annotations (version 91) that included genes with protein coding, lincRNA, macro_lincRNA, immune (IG_*, TR_*), antisense_RNA, and miRNA Ensembl (version 91) biotypes. No normalisation was applied during the aggregation step. Cells with barcodes common to both samples were removed from the analysis to avoid issues associated with index hopping. The aggregated count matrix was randomly down-sampled in order to normalise the median number of UMIs per-cell between the two samples (“downsampleMatrix” function from the DropletUtils R package).

### Identification of sub-populations from single-cell RNA-sequencing data

Cells with <500 genes, >30k UMIs, <0.3% mitochondrial reads, >5% mitochondrial or high expression of *Igkc* and *Igha* genes were removed leaving a total of n=553 WT and n=807 *Irf5*^−/−^ cells. The set of Irf5^−/−^ cells was randomly sub-sampled to match the number of WT cells. Per-cell UMI counts were then normalised, scaled and the effects of total UMI counts, percentage of mitochondrial counts and cell cycle (“all” effects; using known G2 and S phase associated genes^76^) regressed out with the Seurat R package (version 2.3.4). Significantly variable genes (n=1107) were selected using the “trendVar” function from the R Bioconductor package Scran (minimum mean log-expression 0.05, BH adjusted p-value < 0.05). These genes were used as input for principal component analysis (PCA), and significant PCs (n=30) identified using Seurat (“JackStraw” test, BH adjusted p < 0.05). Graph-based clustering of the significant PCs was performed using Seurat (“original” Louvain algorithm, resolution=1.1). Significant cluster markers conserved between the genotypes were identified by intersecting the results of separate tests for cluster markers within each genotype (Seurat “Findmarkers” function, Wilcoxon tests, BH adjusted p value < 0.05). The UMAP projection (Figure 5a) was computed using all of the significant PCs (“RunUMAP” function, Seurat).

### Pseudo-time analysis of single-cell RNA-sequencing data

For pseudo-time analysis the data was first subset to cells identified as Monocytes or Macrophages, and significant variable genes and PCs re-computed as above. The diffusion map (Figure 6a) was computed using all significant PCs (n=20 PCs, “JackStraw” test, BH adjusted p < 0.05) and the R Destiny package^77^. The R package Slingshot (version 0.99.13) was used to fit a minimum spanning tree to the first three dimensions of the diffusion map and to infer the global lineage structure (cluster labels as shown in Figure 5a; starting cluster specified as “Mono1”) ^45^. Slingshot was used to construct a smoothed curve along the single identified lineage and to compute pseudo-time values for each cell. The R package “gam” (version 3.5.0) was used to identify genes significantly differentially expressed across the trajectory (general additive model, BH adjusted p < 0.05).

### Boyden chamber migration assay

Easysep-enriched monocytes were seeded at 20,000 cells/well in 80 μL on, Wells of a 96 well ChemoTX disposable chemotaxis system (Neuro Probe) were filled with 300 μL Boyden Media (RPMI-1640 + 0.1% BSA + 25 mM HEPES) with or without chemotaxis stimuli at desired concentrations. A 96 well, 8 μm pore size mesh was placed on top of filled wells. 80 μL of enriched monocytes at a concentration of 250 cells/μL in Boyden Media was overlain on each membrane well. Cells were incubated for 90 mins at 37 °C, 5% CO_2_ in a humidified HeraCell 150 incubator (ThermoFisher).

After incubation, the media were discarded, and the membrane was rinsed briefly with PBS + 0.1% BSA to remove loosely, and non-adherent monocytes. The membrane was then fixed for 3 hours in PBS + 4% paraformaldehyde solution, rinsed with PBS, and incubated for 3 hrs in PBS + 30% sucrose. The membrane was rinsed in PBS one final time, before air drying. Air-dried membranes were excised from the frame, and experimental groups were mounted on positively charged polysine slides in Glycerol Mounting Medium with DAPI + DABCO (Abcam). Slides were cover slipped, and nail polish was painted around the edges to seal the slides. Slides were kept overnight in the dark at room temperature to air dry.

Slides were then imaged using a BX51 fluorescent microscope (Olympus). Four images/well were acquired at 10X magnification and resolution of 1240 × 1024 pixels. Numbers of cells/field were counted using ImageJ ^78^. For counting, images were converted into 16 bit format. Cells were defined as having a circularity coefficient ≥ 0.8 and between 25 and 400 pixels in diameter.

### Statistical analysis

Data were analysed using Prism V.7 (GraphPad). Statistical tests were performed as indicated in figure legends. Two-sided testing was used in all instances unless indicated.

Differences were considered statistically significant when p ≤ 0.05.

## Data availability

Next generation sequencing datasets are available via the Gene Expression Omnibus (GEO) via accession codes GSE129354 (GM-DMDM data) and GSE129258 (MBMC small bulk and single-cell data).

## Acknowledgements

We thank the High-Throughput Genomics Group at the Wellcome Trust Centre for Human Genetics (funded by Wellcome Trust grant reference 203141/Z/16/Z) for the generation of the sequencing data. We are grateful to Sarah Teichmann (Wellcome Sanger Institute) for help in establishing Smart-Seq2 protocol and generating preliminary data that inspired our subsequent single cell analysis. We would also like to thank Naveed Akbar (University of Oxford) for consultations on the cell migration assay. This work was supported by the Kennedy Trust for Rheumatology Research (ALC, MGV, TEK, DB, SNS), the MRC CGAT programme (SNS), the Versus Arthritis (PhD studentship 209966 to HA), the Novo Nordisk Foundation (Tripartite Immunometabolism Consortium - grant NNF15CC0018486 to IAU), and the Wellcome Trust (Investigator Award 095688/Z/11/Z to FP and 209422/Z/17/Z to IAU).

---

**Supplementary Figure S1:**
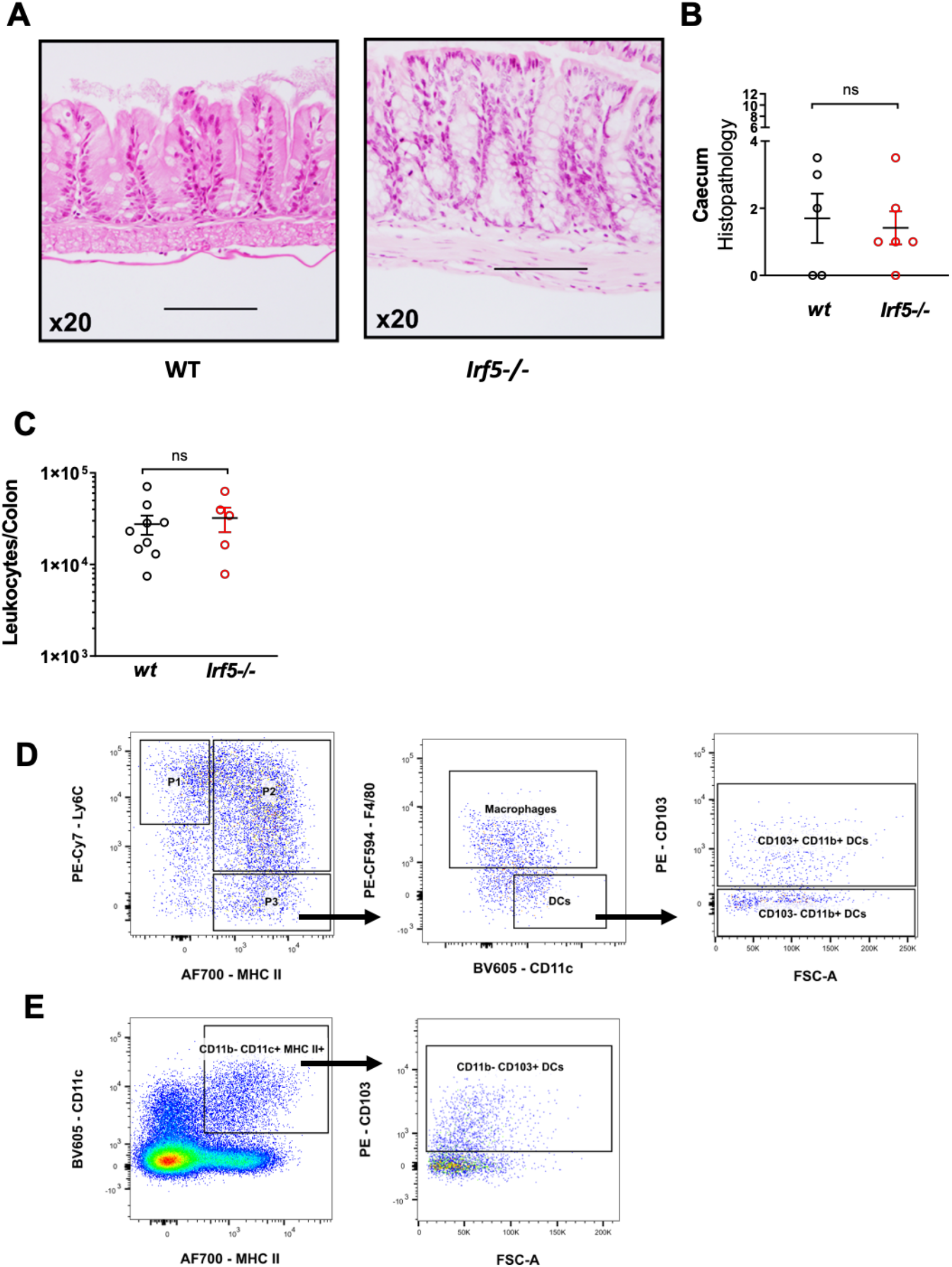
**A)** H&E sections of WT (n=5) and *Irf5*^−/−^ (n=5) caecum at steady state. **B)** Histopathology scoring of steady state caeca. **C)** Number of leukocytes recovered from steady state colons (WT n=9, *Irf5*^−/−^ n=5). **D)** Gating strategy to identify P1 monocytes, P2 monocytes, macrophages, and CD11b^+^ CD103^+^ DCs and CD11b^+^ CD103^−^ DCs. **E)** Gating strategy for CD11b^−^ CD103^+^ DCs. Results are representative of two independent experiments. **B, C)** Mann-Whitney U test. **D)** Gated on LiveCD45^+^Dump^−^CD11b^+^Ly6G^−^SiglecF^−^. **E)** Gated on LiveCD45^+^Dump^−^ CD11b^−^ Data presented are mean ± SEM, ns = not significant

**Supplementary Figure S2:**
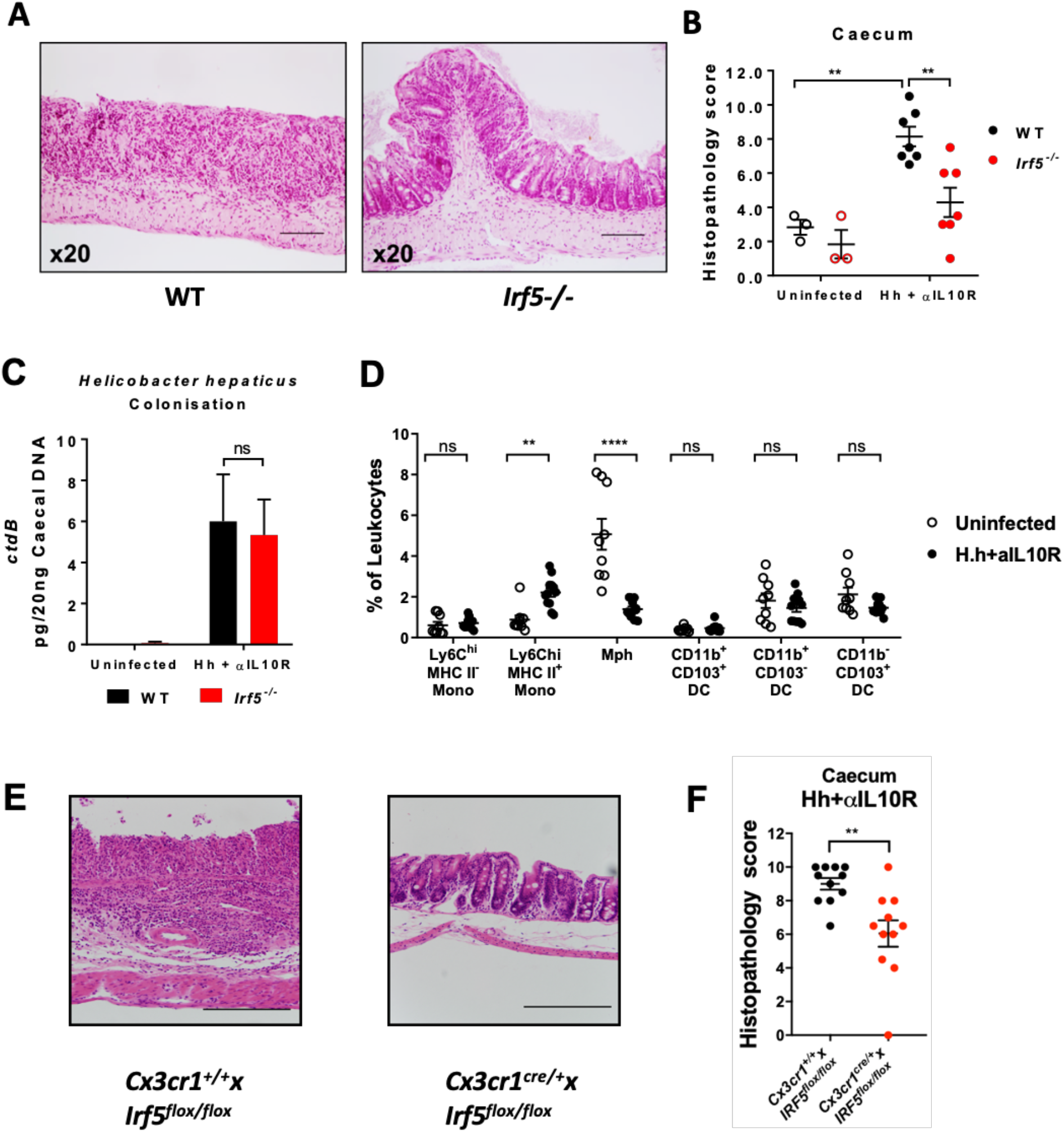
**A)** Representative H&E-stained sections of WT and *Irf5*^−/−^ caecum at d21 Hh + αIL10R colitis. **B)** Histopathology scoring of inflamed caeca (WT ss n= 3 and Hh n =7, *Irf5*^−/−^ ss n= 3 and Hh n =7). **C)** Quantification of *Helicobacter hepaticus* load at d21 colitis (WT ss n= 3 and Hh n =7, *Irf5*^−/−^ ss n= 3 and Hh n =7. **D)** Characterisation of the inflammatory milieu at d21 colitis in WT mice (WT ss n= 9 and Hh n =12). **E)** Representative H&E stained sections of CX_3_CR1^IRF5+^ or CX_3_CR1^IRF^^5−^ caeca at d21 colitis. **F)** Histopathology scoring of CX_3_CR1^IRF5+^ and CX_3_CR1^IRF^^5−^ mice at d21 Hh + αIL10R colitis (WT n=12, *Irf5*^−/−^ n=12). **B, C:** Results are representative of two independent experiments. **D, F:** Data are pooled from two independent experiments. **B,F)** Two-Way ANOVA with Tukey Correction. **C,D)** Two-Way ANOVA with Sidak Correction. Data presented are mean ± SEM, ns = not significant, * p ≤ 0.05, ** p ≤ 0.01, *** p ≤ 0.001, **** p < 0.0001

**Supplementary Figure S3:**
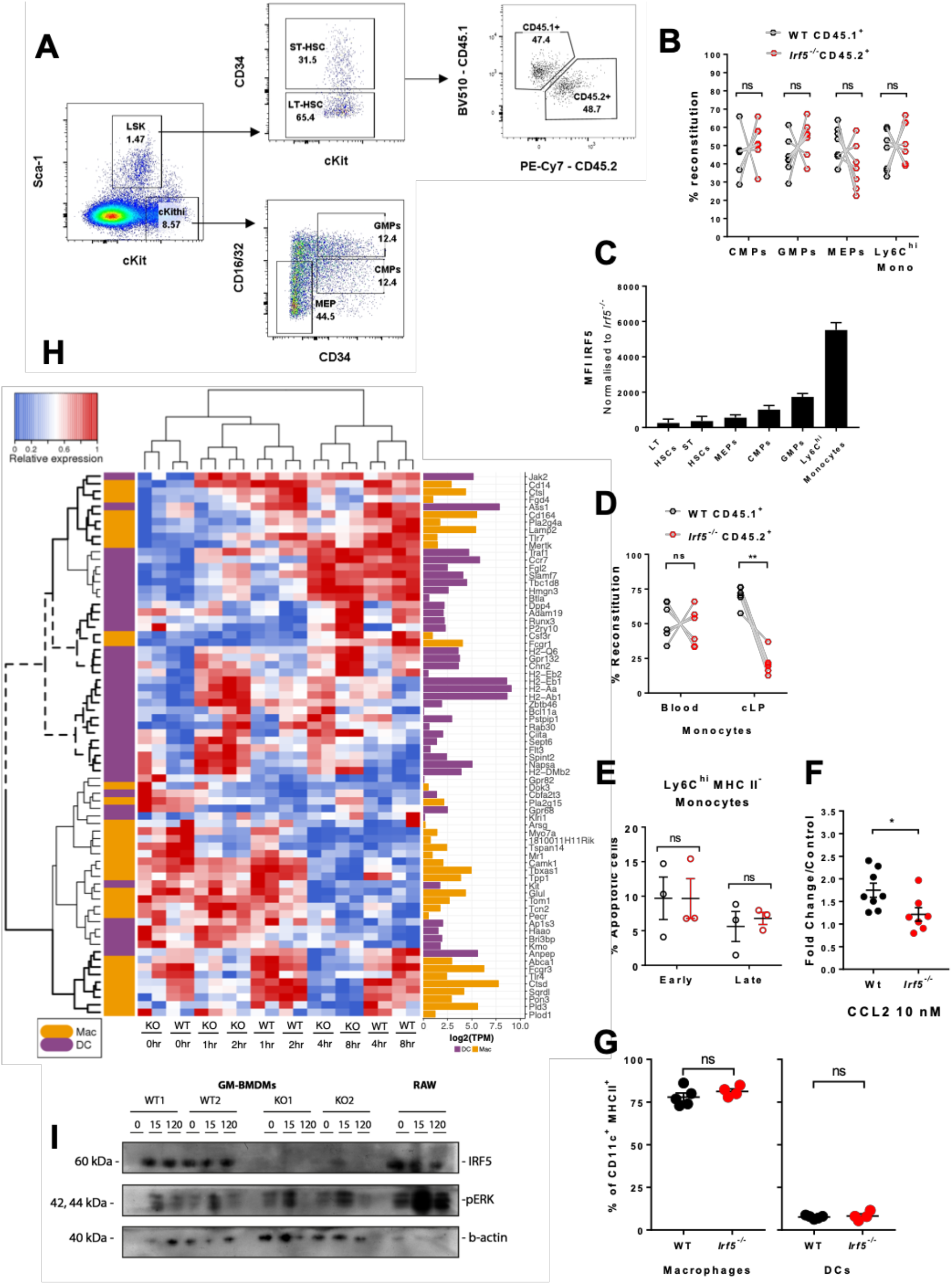
**A)** Gating strategy to identify HSPCs in MBMC. **B)** Quantification of HSPC reconstitution (n=7). **C)** IRF5 expression in the bone marrow quantified by intracellular flow cytometry staining. **D)** Reconstitution of blood and cLP Ly6C^hi^ monocytes (n=7). **E)** Quantification of cell death in Ly6C^hi^ MHC II^−^ (P1) monocytes by flow cytometry using Annexin V and Viability marker staining (n=3). **F)** Boyden chambers were used to quantify the migration potential of isolated splenic monocytes towards CCL2 (n=8). **G)** Flow cytometry analysis of GM-CSF-cultured mouse bone marrow cells (WT n=5, *Irf5*^−/−^ n=4). **H)** Heat map shows mRNA-seq expression of Immgen signature genes for dendritic cells (DC, purple) and macrophages (Mac, yellow) in GM-BMDMs. Expression data are log2(transcripts per million, TPM) scaled per gene, individual replicates are shown. Bold solid dendrogram leaves indicate clusters with increased gene expression in WT, bold dashed dendrogram leaves indicate clusters with increased expression in *Irf5*^−/−^. Bar chart of average expression (log2 TPM) in WT unstimulated cells is shown on the right (WT n=2, *Irf5*^−/−^ n=2, 1, experiment performed once). **I)** IRF5 not required for LPS-induced MAPK activation in macrophages. GM-BMDMs (WT and *Irf5*^−/−^) and RAW264.7 cells were stimulated with 100 ng/ml and 500 ng/ml LPS, respectively at indicated time points. Levels of IRF5, pERK, and b-actin was assessed by Western blot. B, D, G: Data are pooled from two independent experiments. **F:** four pooled experiments. **B, D, E)** Two-Way ANOVA with Sidak Correction. **F, G)** Unpaired t-test Data presented are mean ± SEM, ns = not significant, * p ≤ 0.05

**Supplementary Figure S4:**
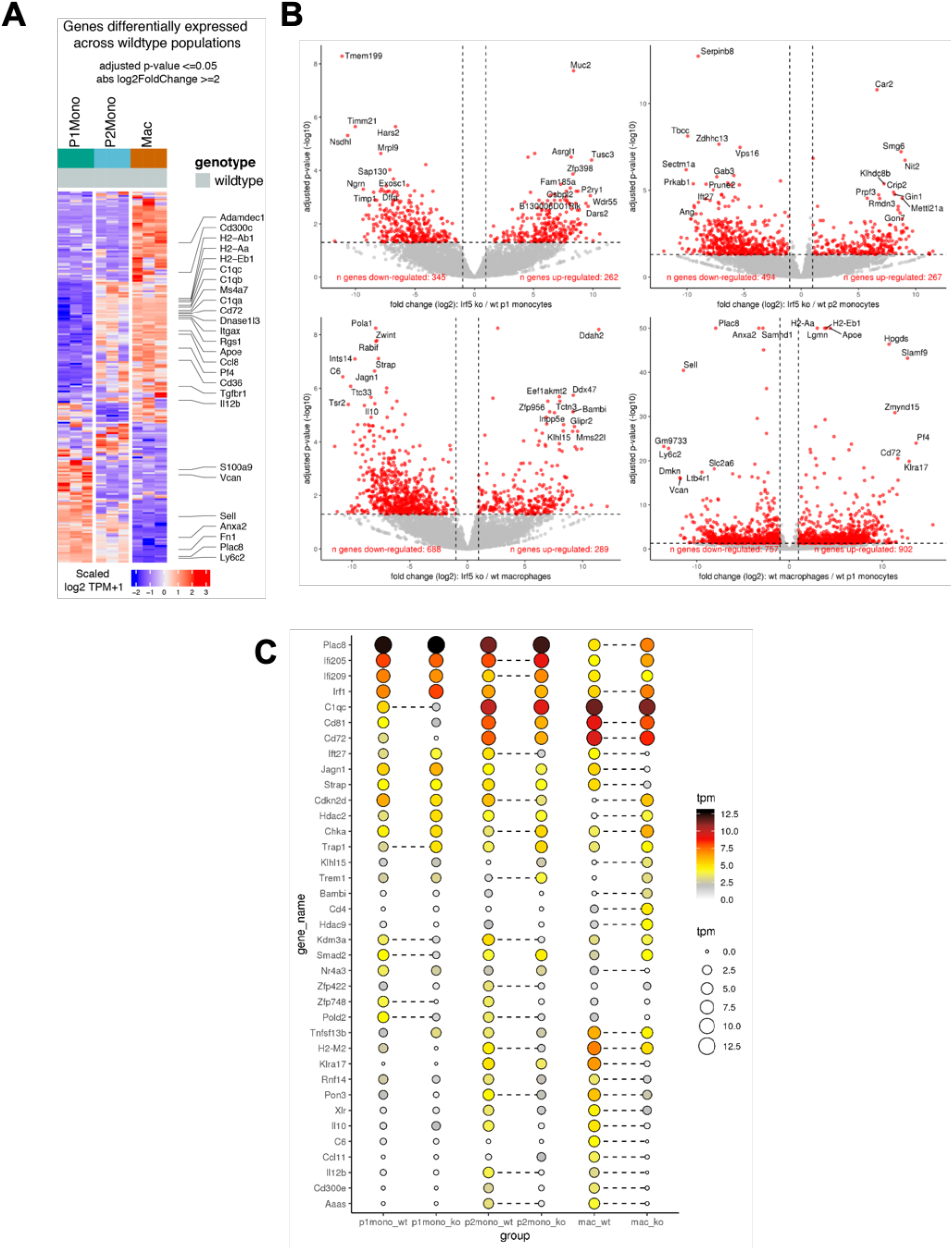
Small-bulk RNA-sequencing analysis of MNP populations during intestinal inflammation (experiment performed once, n=3 chimaeras). **A)** The heatmap shows the expression of genes found to have significant variation in expression between WT P1 monocytes, WT P2 monocytes and WT macrophages isolated from the inflamed cLP of the MBMCs (DESeq2, LRT test, BH adjusted p < 0.05). **B)** The volcano plots show genes found to be significantly differentially expressed (red dots, separate DESeq2 analyses, BH adjusted p < 0.05, | fc | >2) between WT vs *Irf5*^−/−^ P1 monocytes (top left), WT vs *Irf5*^−/−^ P2 monocytes (top right), WT vs *Irf5*^−/−^ macrophages (bottom left), and between WT P1 monocytes vs WT macrophages (bottom right). **C)** The dot plot shows the expression (mean TPM, n=3 biological replicates) of selected genes found to be significantly differentially expressed (dashed lines, | fc | > 2, BH adjusted p < 0.05) between WT and Irf5^−/−^ cells at one or more stages of the monocyte waterfall.

**Supplementary Figure S5:**
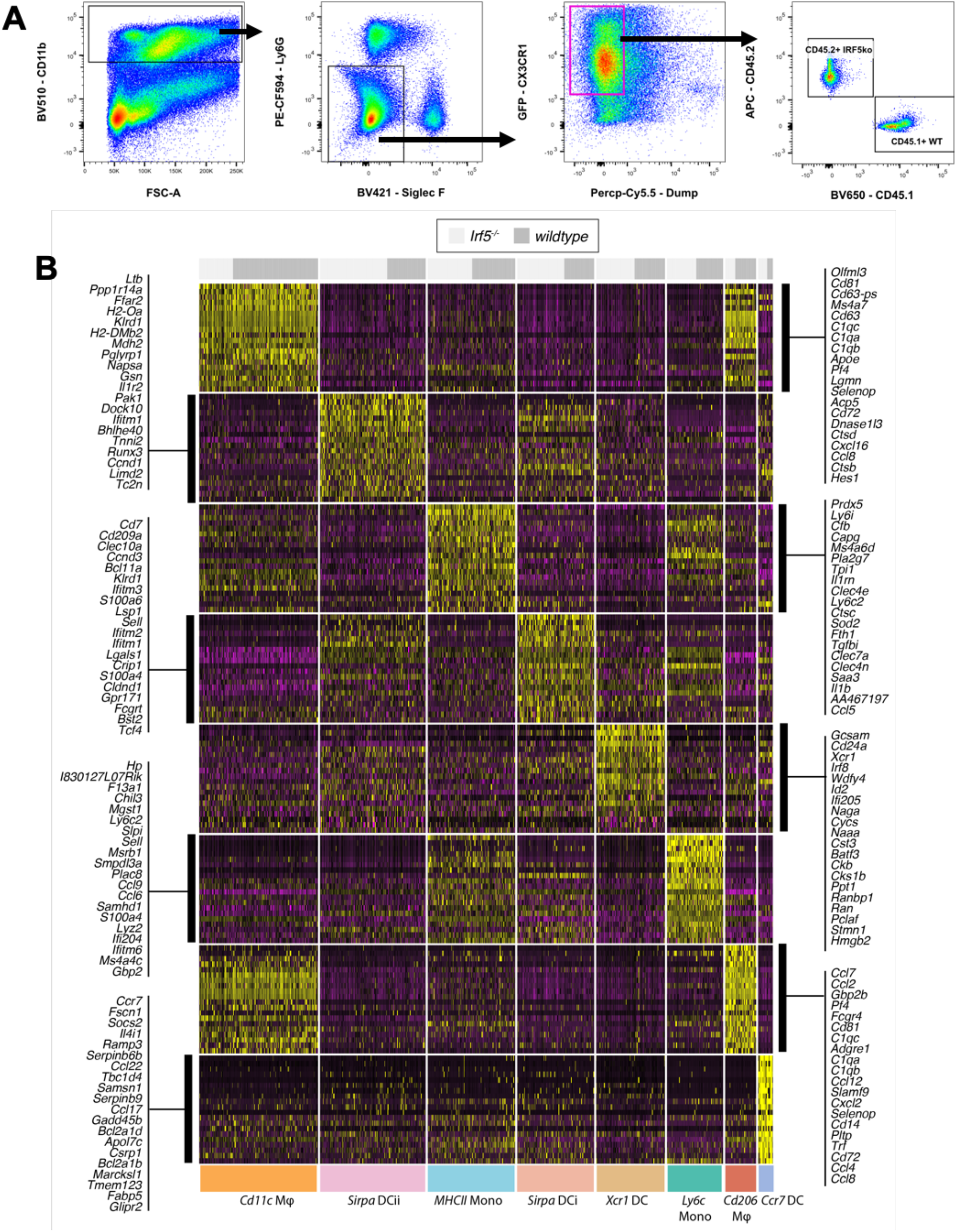
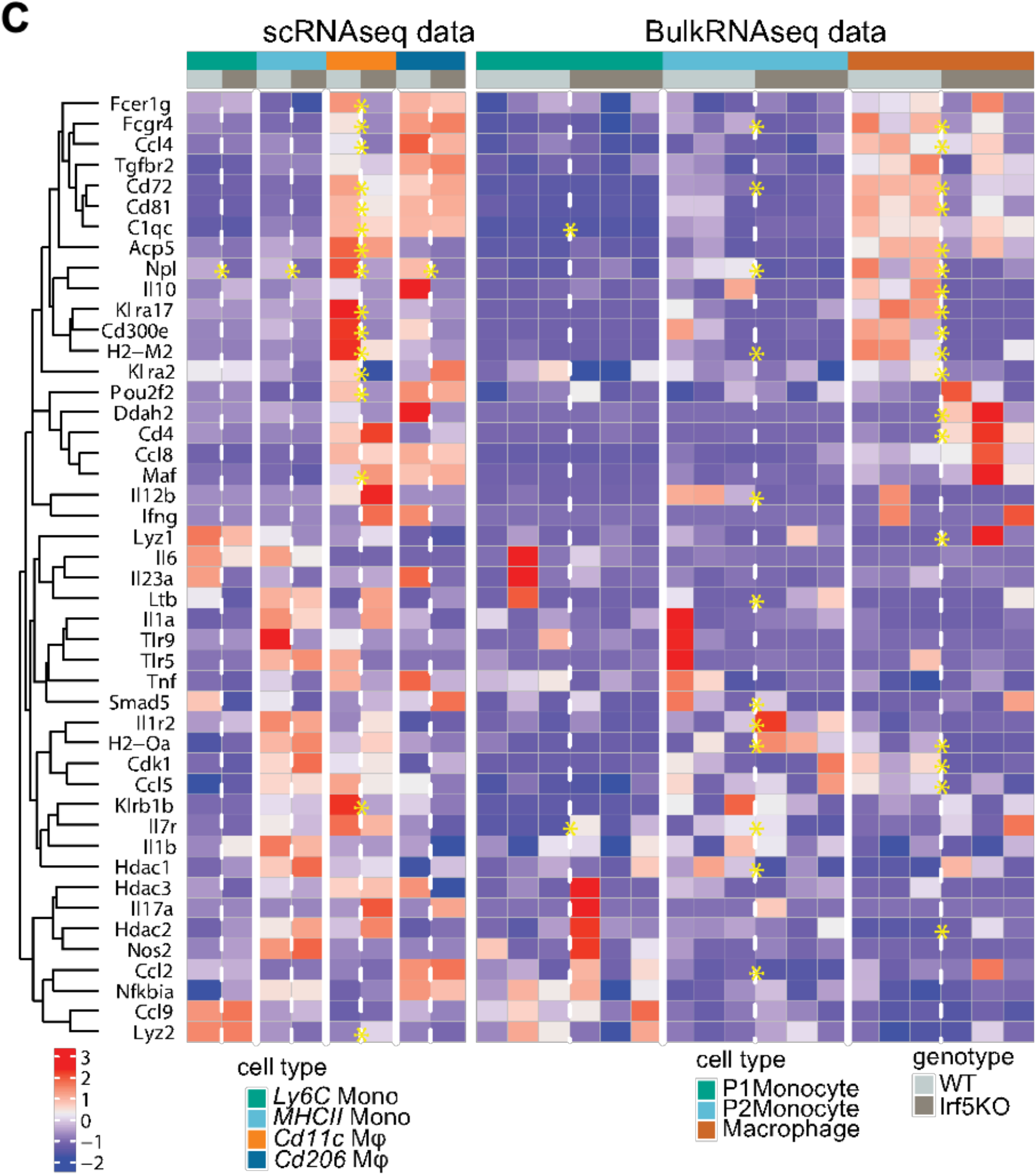
**A)** Representative gating strategy for sorting CX_3_CR1^+^ MNPs from MBMCs, gated on singlets, Live, CD45^+^. **B)** The heatmap shows the expression of the top significant conserved cluster marker genes from the MBMC single-cell RNA-sequencing experiment (Seurat analysis, Wilcoxon tests, BH adjusted p < 0.05 in separate tests of cells of both genotypes). scRNAseq: experiment performed once, n=1 (pool of cLP MNPs from 3 MBMCs) bulk RNAseq: experiment performed once, n=3 chimaeras **C)** The heatmap shows expression of genes found to be significantly differentially expressed (yellow stars, BH adjusted p < 0.05) between WT and *Irf5*^−/−^ MNPs in the single-cell (Wilcoxon tests) or small bulk (DESeq2 analyses) RNA-sequencing datasets.

**Supplementary Figure S6:**
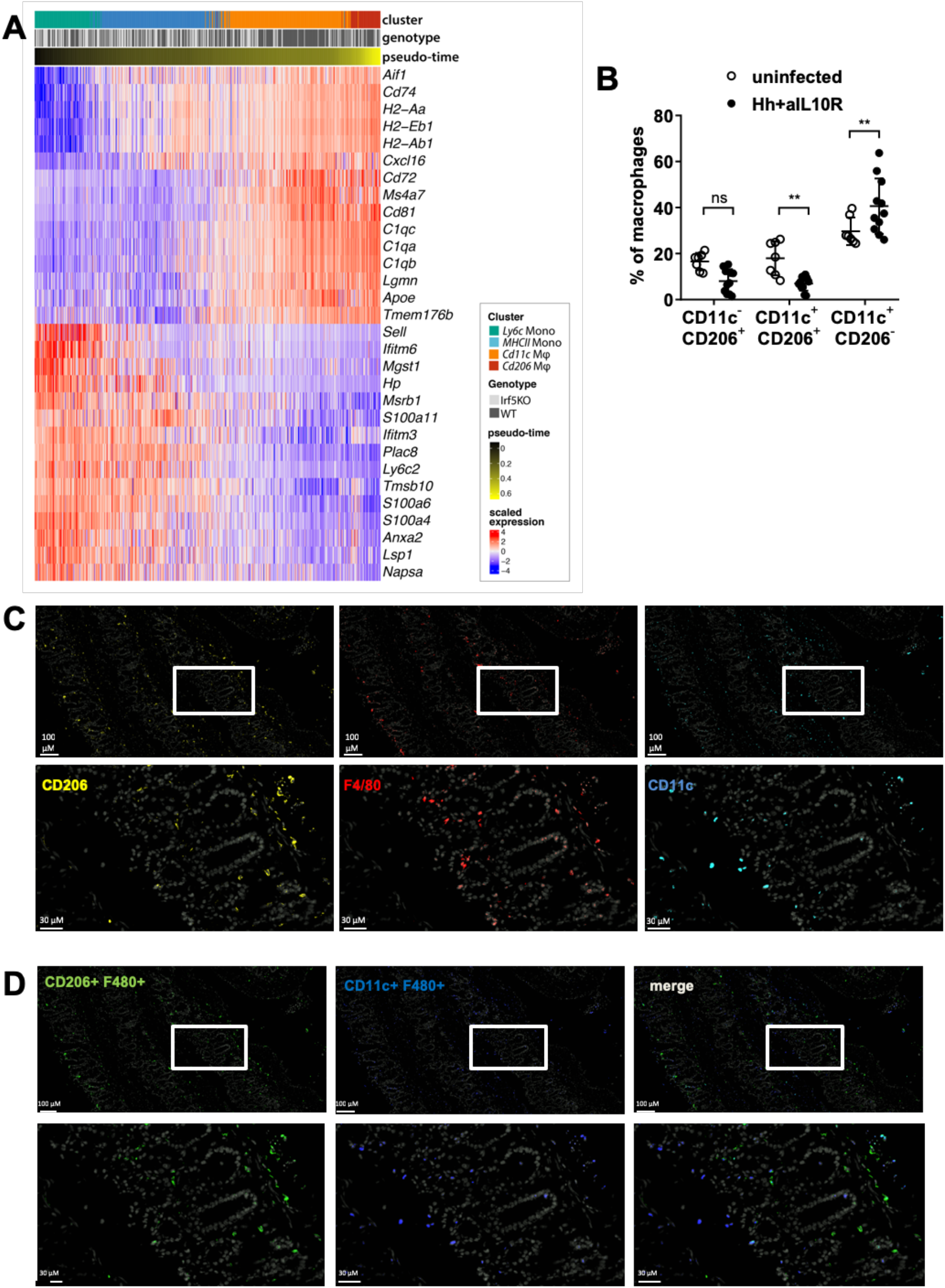
**A)** The heatmap shows the expression of the top thirty genes found to have significant changes (n=1,907 in total) in expression along a smooth curve fitted to the identified pseudo-time trajectory (generalised additive model, BH adjusted p < 0.05). Cells (columns) are ordered by pseudo-time. **B)** The frequency of WT macrophages expressing CD11c or CD206 or Both at steady state (n=7) and at d21 Hh + αIL10R colitis (n=11). **C)** Single channel images of Fig.6C (n=4). **D)** Double-labelled (CD206^+^F4/80^+^, Green, and CD11c^+^F4/80^+^, Blue) macrophage localisation in the steady state mouse cLP by immunohistofluorescence (n=4). **B:** Data are pooled from two independent experiments. **B)** Two-Way ANOVA with Sidak Correction. Data presented are mean ± SEM, ns = not significant, ** p ≤ 0.01

**Supplementary Figure S7:**
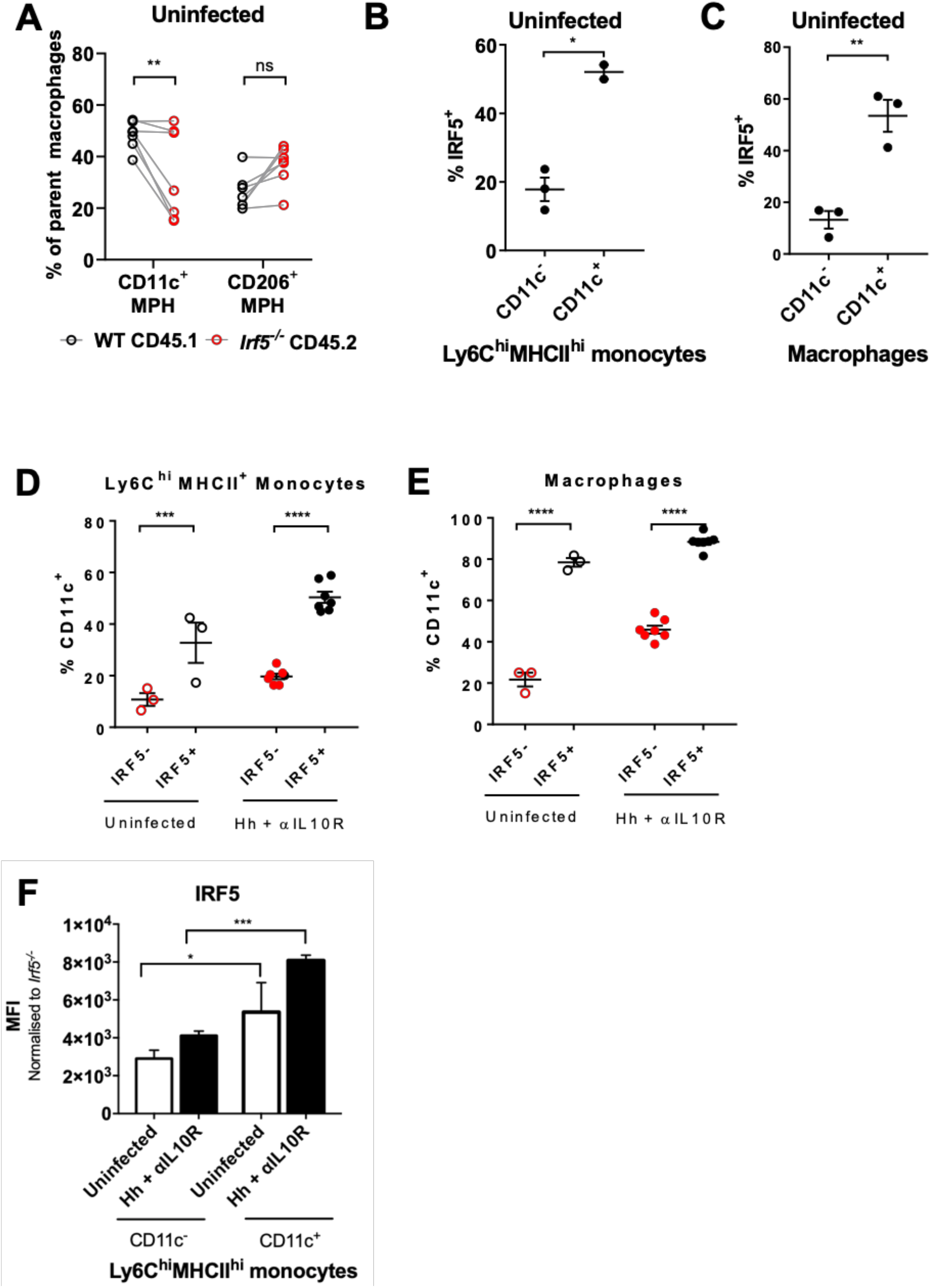
**A)** Frequency of parent macrophages (WT or *Irf5*^−/−^) expressing either CD11c or CD206 in uninfected MBMC (n=7). The percentage of IRF5^+^ CD11c^−^ or CD11c^+^ **B)** P2 monocytes or **C)** macrophages in steady state WT mice (n=3). Frequency of CD11c^+^ cells within IRF5^−^ of IRF5^+^ **D)** P2 monocytes, and **E)** macrophages in the cLP (ss n=3, Hh n=7). **F)** IRF5 expression in CD11c^+^ vs CD11c^−^ P2 monocytes (n=3). **A:** data are pooled from two independent experiments. **B, C, D, E, F:** Data are representative of two independent experiments. **A,D,E)** Two-Way ANOVA with Sidak Correction. **B,C)** Paired t-test. **F)** Two-Way ANOVA with Tukey Correction. Data presented are mean ± SEM, ns = not significant, * p ≤ 0.05, ** p ≤ 0.01, *** p ≤ 0.001, **** p < 0.0001

